# Endocrine-Adapted Pituitary Macrophages Regulate Gonadotropin Secretion through CXCL5-CXCR2-MAPK Signaling

**DOI:** 10.64898/2026.08.12.744557

**Authors:** Zena D. Del Mundo, Jocelyn Ha, Lily Zhou, Antonia Zhang, Gabriela De Robles, Kiara Wiggins, Kien Pham, Naveena Ujagar, Julio Ayala Angulo, Karen Tonsfeldt, Stephanie Correa, Ed Van Veen, Dorota Skowronska-Krawczyk, Dequina A. Nicholas

## Abstract

Chronic inflammation disrupts hormonal balance in the Hypothalamic-Pituitary-Gonadal (HPG) axis, contributing to reproductive disorders. While immune cells in the hypothalamus and ovaries have been extensively studied, their impact on the pituitary remains largely unexplored. Our research identifies pituitary macrophages (PitMacs) as the dominant pituitary immune cell population with a role in regulating reproductive gonadotropin secretion both *in vitro* and *in vivo*. Using a targeted AAV-based depletion strategy, we demonstrate that a reduction of PitMacs decreases serum gonadotropins, luteinizing hormone (LH) and follicle-stimulating hormone (FSH), in female mice. PitMacs are transcriptomically distinct from other tissue-resident macrophages and harbor a unique translational program that reflects the pituitary’s endocrine identity, including active translation of growth hormone (*Gh*) and prolactin (*Prl*). Cytokine profiling identified CXCL5 and IFN-γ as key PitMac-derived mediators of gonadotropin regulation. Mechanistically, CXCL5 signals through CXCR2 to activate the MAPK pathway, converging with Gonadotropin-Releasing Hormone (GnRH) signaling in a time-dependent manner to regulate LH secretion and GnRH receptor surface expression. These findings establish PitMacs as essential endocrine-immune integrators, opening new avenues for understanding inflammation-driven reproductive disorders.

**One Sentence Summary:** Pituitary macrophages are unique hormone-producing immune cells that regulate hormone secretion via cytokine signaling.

## INTRODUCTION

Macrophages are essential immune cells with distinct functions tailored to meet the specific needs of their microenvironment *(1)*. For example, liver macrophages called Kupffer cells clear foreign pathogens and dead red blood cells *(2)*. Macrophages in the lung, or alveolar macrophages, maintain respiratory health by removing inhaled debris and clearing infectious particles such as bacteria and viruses *(3)*. Tissue-specific functions of macrophages extend to the skin, spleen, lymph nodes, muscle, and the brain. A yolk-sac-derived population of resident macrophages was recently discovered in the pituitary, though the tissue-specific function of these cells is unclear *(4)*.

The pituitary gland is the master regulator of the endocrine system, coordinating nearly every physiological process through the secretion of pituitary hormones. These include growth hormone (GH), prolactin (PRL), and the reproductive gonadotropins (luteinizing hormone [LH] and follicle-stimulating hormone [FSH]). Disruption of pituitary hormone secretion underlies a broad spectrum of diseases that collectively affect hundreds of millions of people worldwide. Among physiology governed by the pituitary, the hypothalamic-pituitary-gonadal (HPG) axis orchestrates the reproductive system through hormonal signaling and feedback mechanisms. Disruptions at any level of the HPG axis can cause hormonal imbalance and lead to reproductive disorders such as Polyendocrine Metabolic Ovarian Syndrome (PMOS), endometriosis, and uterine fibrosis, underscoring the importance of understanding the mechanisms governing hormone regulation *(5-8)*.

Inflammation is a significant disruptor of the HPG axis and hormonal homeostasis. Inflammation influences gonadotropin releasing hormone (GnRH) secretion in the hypothalamus through direct cytokine action and modulating estradiol feedback on GnRH neurons *(9, 10)*. In the gonads, lipopolysaccharide (LPS)-induced inflammation disrupts ovarian function by reducing estradiol and progesterone levels, impairing follicular development, and inhibiting ovulation, thereby contributing to cyst formation *(11-14)*. In the pituitary, inflammatory mediators have been shown to suppress or enhance gonadotropin secretion depending on the context. Notably, LPS-induced inflammation affects LH and FSH secretion in a species and concentration-specific manner. Increased LH levels are observed in response to chronic LPS in mice and geese, whereas the opposite occurs in ewes *(13, 15-22)*.

Tissue-resident macrophages are known for their heterogeneity, often having inflammatory activities that suit their microenvironment and tissue function. For example, macrophages in the reproductive tissues contribute to fetal tolerance and disease progression *(23-25)*. In the pregnant uterus, macrophages adopt an anti-inflammatory phenotype to prevent fetal rejection, whereas during labor, they shift to a pro-inflammatory state to facilitate childbirth *(23, 24)*. In pathological conditions such as endometriosis, uterine macrophages contribute to disease progression by promoting inflammation and aberrant tissue growth *(26, 27)*. Despite increasing recognition of the interplay between inflammation and the HPG axis, the tissue-specific function of macrophages in the pituitary is yet to be examined.

In this study, we demonstrate that pituitary macrophages exhibit tissue-specific functions, including production and regulation of gonadotropins, and that secretion of inflammatory cytokines directly regulates gonadotropin secretion. This work provides new insights into the immune regulation of the HPG axis and its implications for reproductive health. Further, these findings establish a foundation for delineating the mechanisms underlying pituitary inflammation and its contribution to inflammation-related hormonal disorders, thereby bridging immunology and pituitary biology.

## RESULTS

### PitMacs regulate hormone secretion *in vitro*

To determine whether pituitary immune cells exert functional consequences on pituitary hormone production, we first examined the correlation between pituitary CD45 expression and LH levels (**Fig. S1A**) *(28)*. Pituitary CD45 expression trended towards an elevation during the LH surge, suggesting that immune cells may be dynamically regulated alongside gonadotrope activity. Therefore, we identified the specific immune cell populations contributing to the bulk of pituitary CD45 expression by profiling pituitary immune cells via single-cell RNA sequencing (scRNA-seq) in healthy female and male mice. This analysis revealed that pituitary macrophages, hereafter referred to as PitMacs, constitute the predominant immune cell population in the pituitary gland (**Fig. 1A-B**), a finding independently validated by flow cytometry (**Fig. 1C, S1B**) *(4)*.

**Fig. 1.**
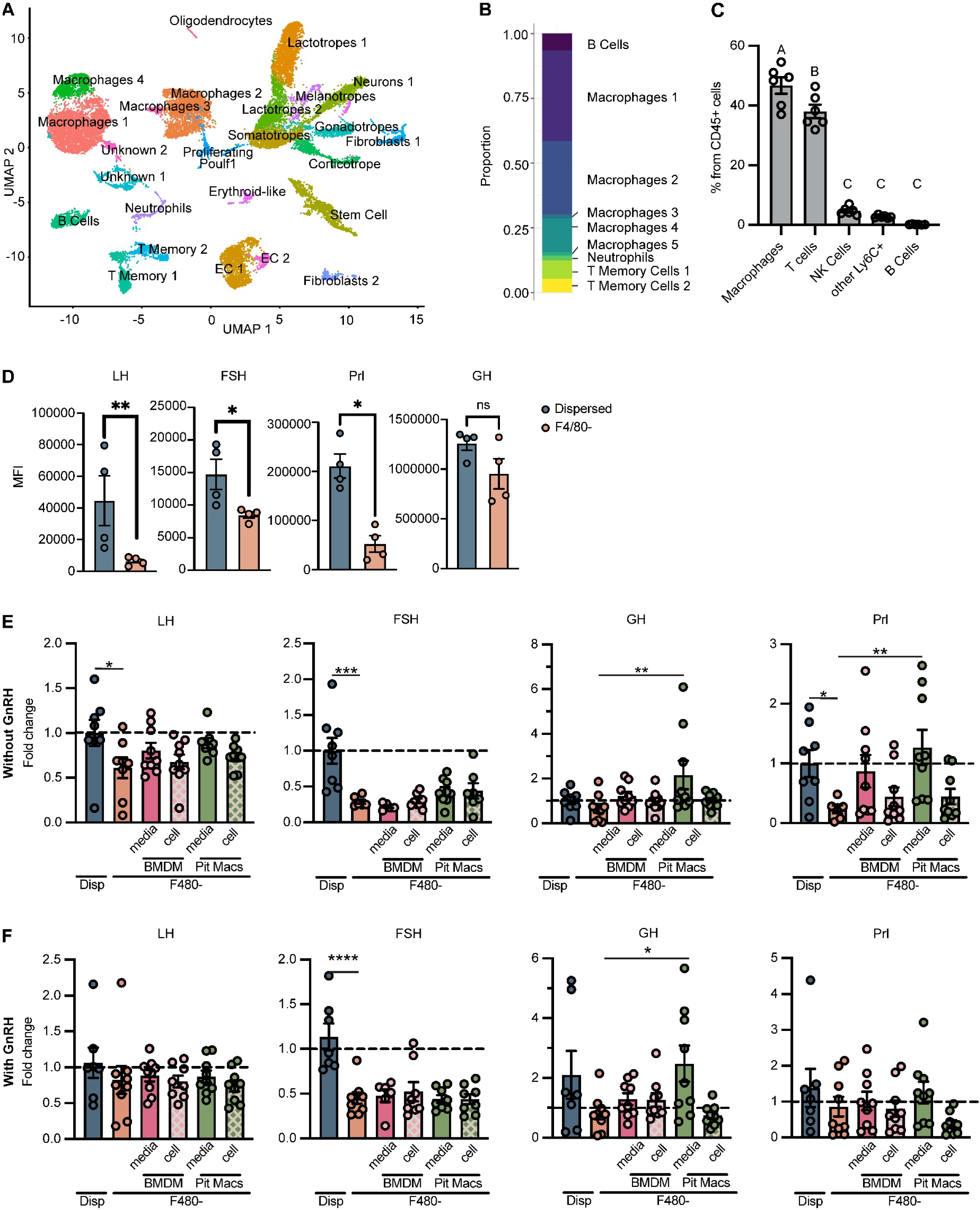
PitMacs regulate hormone secretion *in vitro*. (**A**) Pituitaries from healthy 10-week-old male and female C57BL/6J mice were collected for single-cell RNA sequencing (scRNA-seq). CD45+ immune cells were enriched using positive-selection magnetic beads prior to sequencing. (**B**) Bar graph showing the proportion of pituitary immune cell clusters identified in scRNA-seq data from healthy male and female C57BL/6J mice. (**C**) Immune cell populations were validated using a spectral flow cytometry panel. (**D**) Culture supernatants from dispersed or macrophage (F4/80)-depleted pituitaries (n = 5) were analyzed by Luminex to quantify LH and FSH levels, reported as mean fluorescence intensity (MFI). Statistical comparisons were performed using a Student’s t-test, *p <0.05, **p <0.01. (**E-F**) Macrophage-depleted primary mouse pituitaries were rescued *in vitro* with either PitMacs, bone marrow-derived macrophages (BMDMs), or their respective 12-hour conditioned media at a ratio of 2 gonadotropes per 1 macrophage. For GnRH-stimulated conditions (F), cultures were treated with 10 nM GnRH for 30 minutes prior to supernatant collection. Hormone levels were quantified by Luminex. Outliers were removed prior to Box-Cox transformation, and statistical comparisons were performed using Dunnett’s Test relative to F4/80 depleted groups, *p <0.05, **p <0.01, ***p <0.001, ****p <0.0001.

Given the relevance of pituitary function to reproductive hormone disorders that are common in women, subsequent studies were focused on female mice. To determine the extent to which PitMacs contribute to neuroendocrine control of gonadotropin secretion, we depleted PitMacs from mouse pituitaries *in vitro* using an F4/80 positive selection kit. We show that depletion of PitMacs results in a significant loss of LH and FSH secretion in GnRH-treated conditions (**Fig. 1D**). We also found that PitMac depletion reduces prolactin (Prl) but not growth hormone (Gh) (**Fig. 1D**).

To identify the mechanism underlying the observed reduction in pituitary hormone levels following macrophage depletion, we investigated canonical macrophage functions and classified them into two broad categories, direct and indirect regulation. We sought to determine whether macrophage-mediated control of pituitary hormone secretion operates through direct mechanisms such as paracrine cytokine secretion or cell-to-cell contact with gonadotropes, or through indirect mechanisms, including downstream cytokine signaling cascades and immune cell activation. To this end, we co-cultured PitMac-depleted pituitaries with either primary PitMacs or PitMac-conditioned medium. To further assess whether the observed effects on pituitary hormone secretion are specific to PitMacs or shared by other macrophage populations, we included bone-marrow-derived macrophages (BMDMs) and their conditioned media as additional treatment groups.

In non-GnRH stimulated pituitaries, both PitMacs and BMDMs and their conditioned media partially rescued LH levels that were reduced upon macrophage depletion. GH and PRL trended toward a decrease following macrophage depletion, and addition of PitMac conditioned media elevated both hormones above depleted levels (**Fig. 1E**). Similar trends were observed in GnRH-stimulated pituitaries, though these did not reach statistical significance for LH and PRL (**Fig. 1F**). GH, in contrast, showed a consistent response from PitMac-conditioned media in both conditions (**Fig. 1E-F**). Together, these findings suggest that soluble factors secreted by macrophages contribute to the regulation of pituitary hormone secretion in a GnRH-and macrophage identity-dependent manner.

### PitMacs have a unique transcriptional and functional profile compared to other tissue resident macrophages

To extend our *in vitro* findings to an animal model with an intact HPG axis, we sought to develop a pituitary-specific *in vivo* macrophage depletion strategy that would selectively target PitMacs without depleting other tissue-resident macrophage populations. To achieve this specificity, we first identified markers uniquely enriched in PitMacs relative to macrophages from other tissues. We integrated our scRNA-seq dataset from healthy male and female pituitaries with the publicly available Tabula Muris dataset and compared immune cell subsets across multiple tissues, including brain, adipose tissue, and liver, following standard preprocessing and batch correction *(29)*. Importantly, differential expression analysis distinguished PitMacs from microglia through the canonical markers *Tmem119*, *P2ry12*, and *Cx3cr1* (**Fig. 2A-B, S2A**). PitMacs were also transcriptomically distinct from, other tissue-resident macrophages, circulating monocytes, and other immune cell populations (**Fig. 2A, S3A**). To further resolve the functional transcriptional identity of pituitary macrophage subsets, differentially expressed genes (DEGs) were filtered against gene lists associated with macrophage activity such as activation, migration, and phagocytosis (**Fig. S4A**). Differential Gene Set Enrichment Analysis (DGSEA) further revealed PitMac subset-specific genes involved in biological activities including interferon activation, heat shock responses, and hormone production (**Fig. S5A-D**). Notably, PitMacs expressed transcripts encoding growth hormone (*Gh*) and prolactin (*Prl*), which are hormones canonically produced by the pituitary endocrine cell types somatotropes and lactotropes, respectively (**Fig. 2B**). This hormone-associated transcriptional program, likely shaped by the pituitary niche, suggested that pituitary-enriched hormone transcripts could serve as candidate markers for the design of a PitMac-specific *in vivo* targeting strategy.

**Fig. 2.**
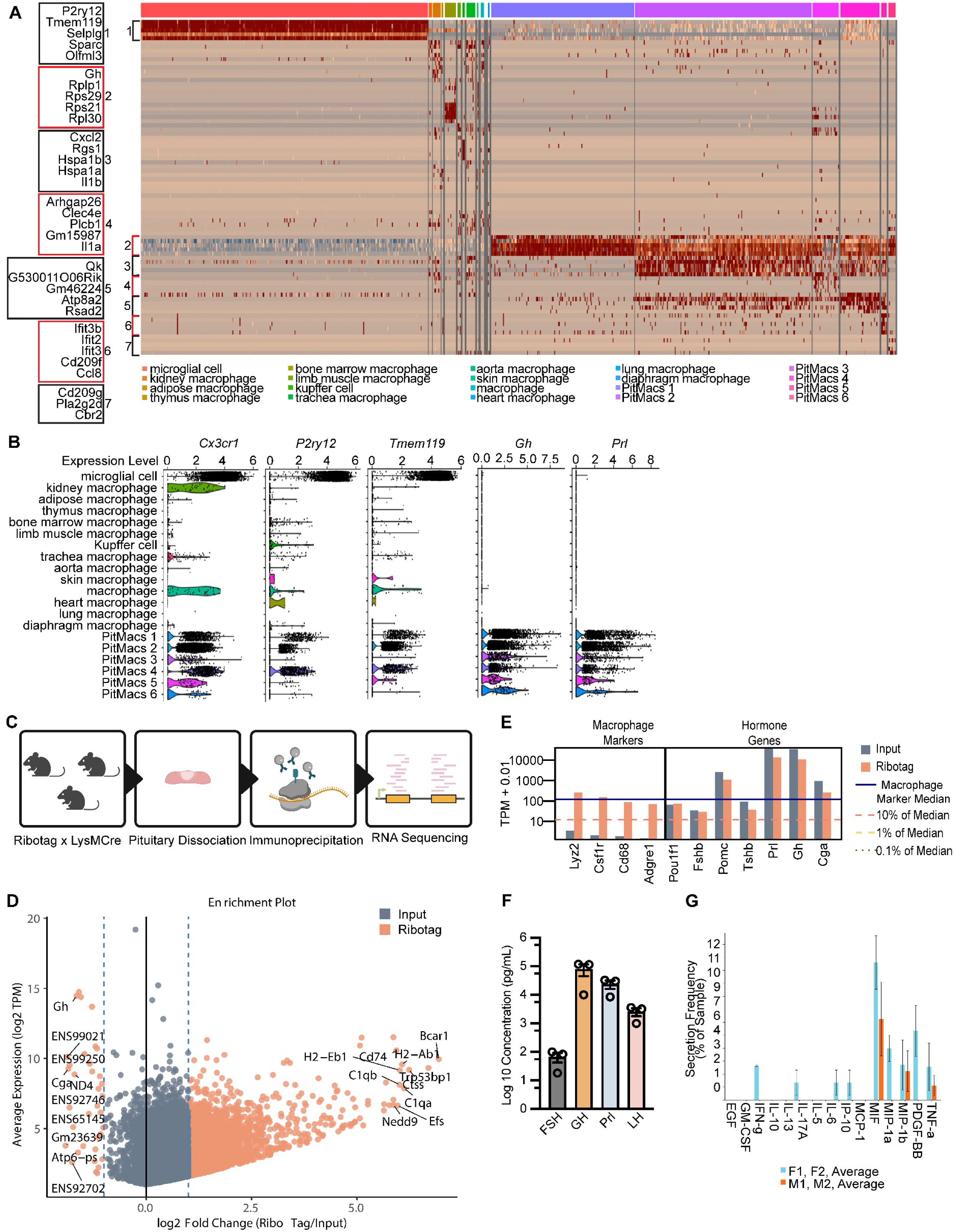
PitMacs have a unique transcriptional and functional profile compared to other tissue resident macrophages. (**A**) Heatmap displaying the top differentially expressed genes (DEGs) in macrophages from distinct tissue depots in the integrated Tabula Muris and pituitary scRNA-seq datasets. The top five DEGs from microglia and PitMacs are labeled. (**B**) Expression levels of microglial markers *Tmem119*, *P2ry12*, and *Cx3cr1*, as well as the genes encoding for hormones GH and Prl across tissue-resident macrophage populations. (**C**) Schematic workflow for isolating actively translated mRNAs from PitMacs. RiboTag mice were crossed with LysMCre mice, and heterozygous offspring were used for pituitary collection. HA-tagged ribosomes from dissociated pituitaries were immunoprecipitated, and associated RNAs were isolated and processed for RNA sequencing. Male and female samples were processed independently. (**D**) Enrichment plot showing genes highly enriched in the immunoprecipitated (RiboTag) fraction relative to total input in female mouse samples. (**E**) Expression levels of canonical macrophage markers (*Lyz2*, *Csf1r*, *Cd68*, and *Adgre1*) are shown alongside genes of interest encoding hormones. Expression values are presented as Transcripts Per Million (TPM), with the percent median TPM of macrophage markers used as a baseline reference. (**F**) PitMacs isolated from 10 week-old female mice were cultured for 12 hours. Hormone secretion was measured using Luminex multiplex assay. Limit of detection values for FSH, GH, Prl, and LH are 6.89, 0.57, 2.20, and 0 pg/mL. (**G**) PitMacs isolated from 10-week-old female and male mice were pre-treated with 10 µg/mL LPS for 24 hours, followed by overnight incubation in LPS-free media to characterize single-cell secretome profiles. Bars are shown as average of n=2 samples from either female (F1,F2) or male (M1,M2) mice.

Given that PitMacs have high expression of phagocytosis-related genes, we next sought to determine whether their hormone transcript expression reflects active translation or passive acquisition of transcripts through phagocytosis of neighboring hormone-producing cells. To address this, we employed a RiboTag approach using LysMCre x HA-RiboTag heterozygous mice, in which HA-tagged ribosomes are selectively expressed in myeloid cells. Pituitaries were dissected and HA-tagged ribosomes were immunoprecipitated from pituitary lysates, enabling selective isolation of mRNAs undergoing active translation in PitMacs (**Fig. 2C**). HA-tagged ribosome immunoprecipitation effectively enriched for macrophage-associated transcripts while depleting markers of non-immune pituitary cell types, confirming the specificity of the approach (**Fig. 2D, S6C**).

To determine whether hormone transcripts in PitMacs were being actively translated at biologically meaningful levels, we used the translational output of canonical macrophage markers as a reference benchmark. Strikingly, *Prl* and *Gh,* hormones typically produced by lactotropes and somatotropes respectively, were translated at levels that exceeded those of canonical macrophage markers in PitMacs (**Fig. 2E**). Additional actively translated transcripts, including other pituitary hormones, hormone receptors, and cytokine receptors, were identified in both female and male mice (**Fig. S6B, S6D-F**). To confirm that PitMacs actively secrete pituitary hormones, PitMacs were isolated from female mice, cultured for 12 hours, and the conditioned media was assessed for hormones using a Luminex multiplex assay. PitMacs secreted significant amounts of LH, FSH, GH, and PRL above baseline detection thresholds (**Fig. 2F**). As additional characterization, we performed a single-cell secretome analysis of PitMacs using a 15 cytokine Isoplexis panel. Here, we show that female and mice exhibit distinct cytokine secretory profiles and polyfunctional strength indices following LPS stimulation, with female PitMacs showing a more diverse cytokine secretion capacity (**Fig. 2G, S6G**). Collectively, these findings reveal that PitMacs harbor a unique translational program that closely reflects the endocrine identity of the pituitary gland, distinguishing them from tissue-resident macrophages in other tissues.

### PitMacs regulate hormone secretion *in vivo* in intact HPG axis

We identified accessible chromatin at the *Gh* locus in PitMacs through secondary analysis of published mouse pituitary ATAC-seq data from Wallis et al. (**Fig. S7A**) *(30-32)*. Based on these data and our finding that PitMacs express hormone transcripts, we selected the *Gh* promoter as a PitMac-specific regulatory element for the design of a targeted *in vivo* depletion construct (**Fig. 2B, 3A**). We developed a pituitary macrophage-depleting AAV using serotype AAV5, which we first validated for efficient pituitary transduction in wild-type mice using an AAV-CMV-GFP construct and subsequently confirmed Cre-dependent expression using an AAV-GFP reporter in LysMCre mice (**Fig. S8**). We then designed a dual Gh promoter-and Cre-dependent AAV construct encoding an auto-cleaving caspase-3, which was delivered via retro-orbital injection into LysMCre mice to selectively induce apoptosis in PitMacs (**Fig. 3A-B**). Three weeks post-injection, PitMacs were depleted by 23.48% on average while lung macrophages remained unaffected, demonstrating the pituitary specificity of this approach (**Fig. 3C-D**).

**Fig. 3.**
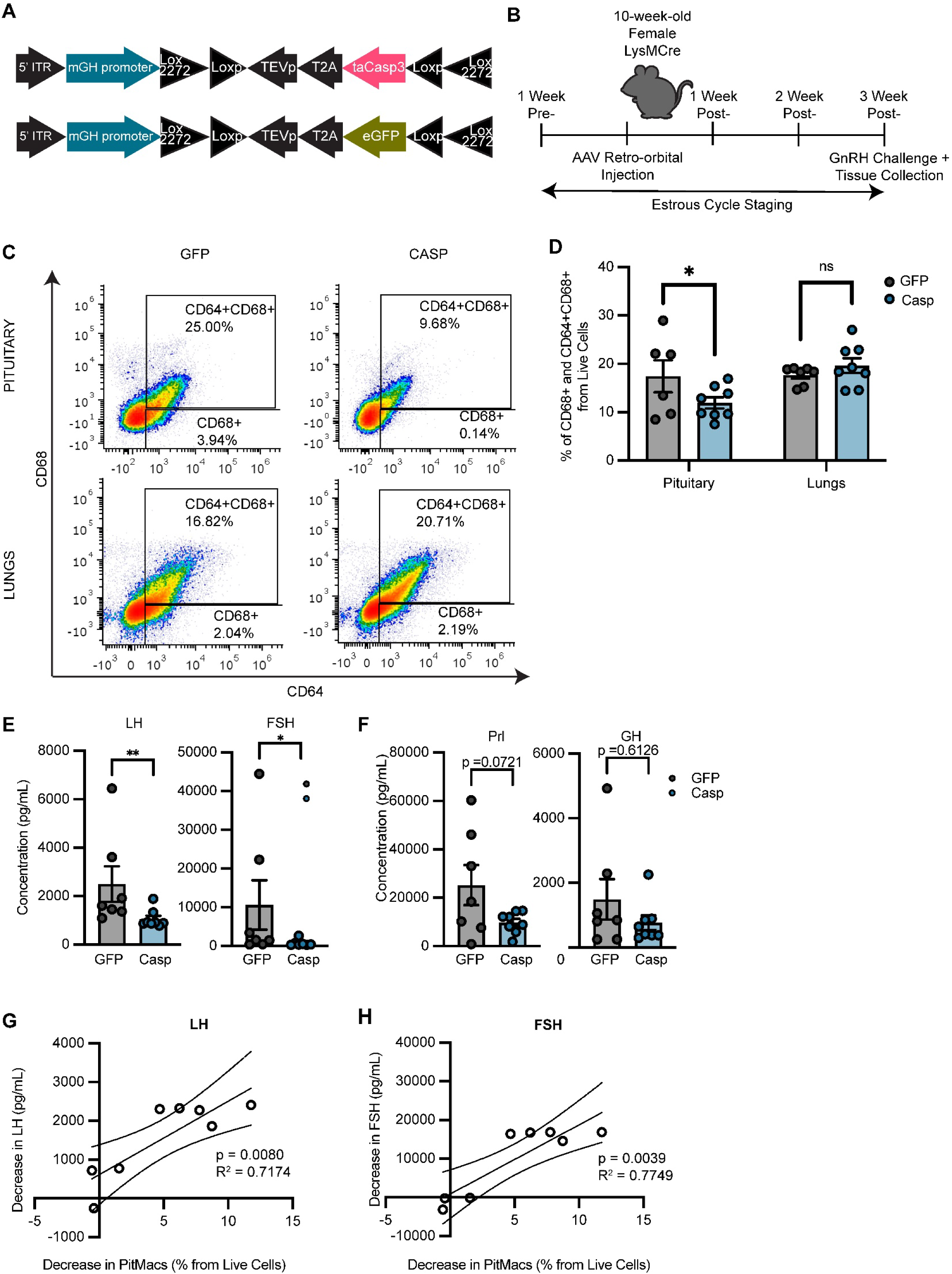
PitMacs regulate hormone secretion *in vivo* in intact HPG axis. (**A**) Schematic of adeno-associated virus (AAV) constructs used for targeted macrophage depletion. An AAV carrying a GH-promoter-driven, Cre-dependent caspase cassette was delivered to LysMCre mice to selectively ablate PitMacs; a Cre-dependent GFP cassette served as a control. (**B**) Experimental workflow where estrous cycle staging was performed 1 week prior to and 3 weeks following retro-orbital AAV injection. GnRH challenge was administered weekly beginning 1-week post-injection to assess serum gonadotropin responses. (**C–D**) Flow cytometry was used to quantify macrophage populations in the pituitary and lungs (peripheral control) 3 weeks post-injection. (**E-F**) Serum LH, FSH, PRL, and GH concentrations (pg/mL) post-GnRH challenge were measured by Luminex. (**G**-**H**) Pearson correlation analysis depicting the magnitude of decrease in serum LH and FSH levels (pg/mL) versus the magnitude of decrease in the PitMac population (% of live cells). Dashed lines represent 95% confidence bands of the best-fit line. (**D** - **F**) Hormone concentrations and macrophage populations were analyzed following appropriate outlier exclusion tests, normality tests, and Box-Cox transformation. Mann-Whitney test was used for LH, FSH, and GH. Unpaired t-test was used for Prl. *p = 0.01–0.05; **p = 0.001–0.01.

Upon pituitary macrophage depletion *in vivo*, we observed a significant decrease in LH and FSH serum levels and a decreasing trend for PRL and GH in response to *in vivo* GnRH challenge (**Fig. 3E-F, S9A**). No changes were observed in the estrous cycle, with mice displaying normal estrous cyclicity, percent time spent in each stage, and estrous cycle length (**Fig. S9B-D**). We also show that the decrease in PitMac population in the pituitary is positively correlated with the magnitude of LH and FSH reduction, but not with PRL and GH levels (**Fig. 3G-H, S9E-F**). The unique role of PitMacs in hormone regulation is further supported by a separate experiment in which whole-body macrophages were depleted using clodronate liposomes, which did not result in gonadotropin level difference (**Fig. S9G**). These findings demonstrate that PitMac regulate gonadotropin production *in vivo*.

### PitMacs modulate hormone secretion through cytokine signaling

Given that macrophages are well-established cytokine-producing cells, we next profiled cytokine secretion in dispersed and macrophage-depleted pituitaries from the *in vitro* macrophage depletion experiment (**Fig. 1D**). Of the 27 cytokines measured, significant reductions in CXCL5 and IFN-γ were observed in association with the decrease in LH, FSH, and PRL (**Fig. 4A-C**). We further demonstrate that the reduction in IFN-γ following macrophage depletion is GnRH-dependent, whereas the reduction in CXCL5 is GnRH-independent (**Fig. 4B, 4C**). This GnRH-independent reduction of CXCL5 suggests that CXCL5 may contribute to gonadotrope regulation in a more constitutive rather than stimulus-dependent manner. Although **Fig. S2F** confirms IFN-γ secretion by LPS-stimulated PitMacs from female mice, macrophages from other tissue compartments are not broadly recognized as producers of CXCL5 or IFN-γ. Furthermore, our single-cell secretome panel did not include CXCL5, and a prior scRNA-seq study identified only IFN-β as a candidate cytokine secreted by PitMacs *(4)*. To address these limitations, we employed a spectral flow cytometry panel to characterize intracellular cytokine expression across PitMac populations. We found that 48.6% of PitMacs are IFNβ^+^IFNγ^−^CXCL5^+^, 32.3% are IFNβ^+^IFNγ^−^CXCL5^-^, 12.1% are IFNβ^+^IFNγ^+^CXCL5^+^, and 0.3% are IFNβ^+^IFNγ^+^CXCL5^-^(**Fig. 4D**).

**Fig. 4.**
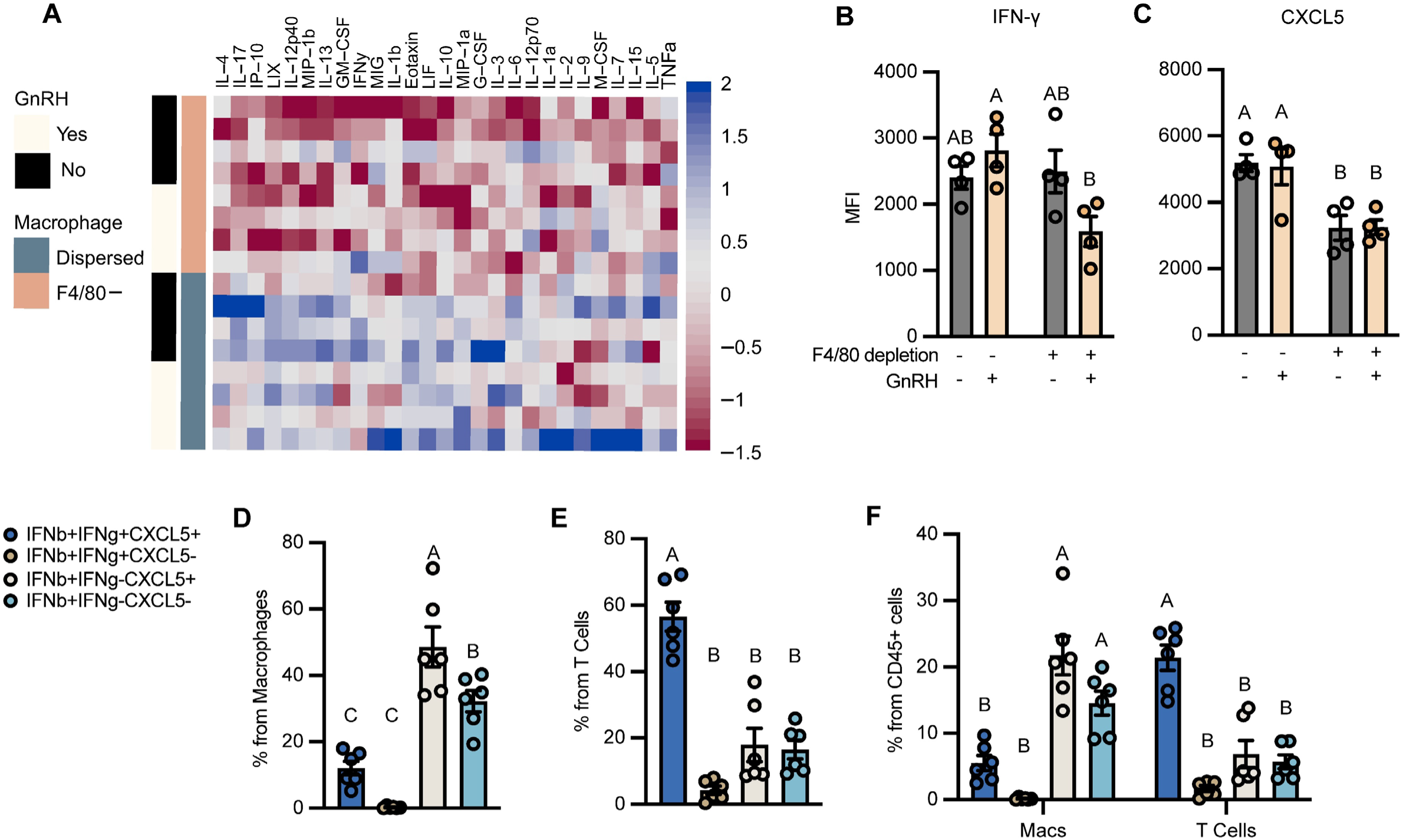
PitMacs modulate hormone secretion through cytokine signaling. (**A**) A 27-plex cytokine Luminex panel was applied to supernatants from the *in vitro* macrophage depletion experiment. A heatmap displays MFI values for 27 cytokines across four conditions: with or without GnRH stimulation and with or without F4/80-mediated macrophage depletion. (**B**-**C**) Bar graphs showing IFNγ (**B**) and CXCL5 (**C**) as the two cytokines significantly depleted following macrophage removal. (**D-F**) Spectral flow cytometry was performed on dissociated pituitaries from 10-week-old female C57BL/6J mice following a 4-hour incubation with a cell activation cocktail (PMA, ionomycin, and Brefeldin A) and intracellular staining. Bar graphs show the percent of population expressing target cytokines within (**D**) macrophages (F4/80+CD11b+), (**E**) T cells (CD3+CD19−), and (**F**) total immune cells (CD45+). Statistical comparisons were performed using Tukey’s HSD test following appropriate outlier removal, normality testing, and Box-Cox transformation. Groups denoted by different letters are statistically significantly different from one another.

To contextualize these findings relative to other pituitary-resident immune cells, we assessed intracellular cytokine expression in pituitary T cells, B cells, NK cells, and Ly6C+ cells (**Fig. 1C**). Significant expression of the target cytokines was detected only in T cells, where 56.6% are IFNβ^+^IFNγ^+^CXCL5^+^, 17.9% are IFNβ^+^IFNγ^-^CXCL5^+^, 16.5% are IFNβ^+^IFNγ^-^CXCL5^-^, and 4.3% are IFNβ^+^IFNγ^+^CXCL5^-^(**Fig. 4E**). While IFNβ expression was shared across PitMacs and T cells, PitMacs represented the predominant single-positive IFNβ+ population (**Fig. 4F**). Importantly the IFNβ^+^IFNγ^-^CXCL5^+^ profile was significantly enriched in PitMacs. Given that IFN-β is expressed across different pituitary immune populations and that CXCL5 shares an overlapping activation pathway with the GnRH receptor (GnRHR), we focused subsequent mechanistic investigations on CXCL5 as a pituitary macrophage-specific mediator of gonadotrope regulation. Together, these findings show that PitMacs are a source of secreted cytokines that may regulate pituitary hormone secretion.

### PitMacs modulate LH secretion through CXCL5-CXCR2 activation

Given the enriched CXCL5+IFN-γ macrophage population observed in the pituitary relative to other immune cell populations, we focused subsequent mechanistic analyses on CXCL5 signaling in gonadotropes. To further dissect the effects of cytokine signaling specifically in gonadotropes, we employed LβT2 cells, a mouse gonadotropic cell line that constitutively produces LH in the absence of exogenous stimulation. We confirmed that LβT2 cells respond to both IFNγ and CXCL5 in a dose-dependent manner (**Fig. S10A, S10B**). As a baseline, we first characterized the effects of CXCL5 and GnRH on LH secretion, GnRHR expression, and CXCR2 expression. At 30 minutes, CXCL5-induced LH secretion trended toward an increase only under non-GnRH treated conditions. In contrast, CXCL5-induced LH secretion at 12 hours was significantly elevated in a GnRH-dependent manner, suggesting that sustained CXCL5 signaling progressively engages GnRH-coupled pathways (**Fig. 5A**). This significant increase in LH secretion at 12 hours was accompanied by a significant upregulation in GnRHR receptor expression (**Fig. 5B**).

**Fig. 5.**
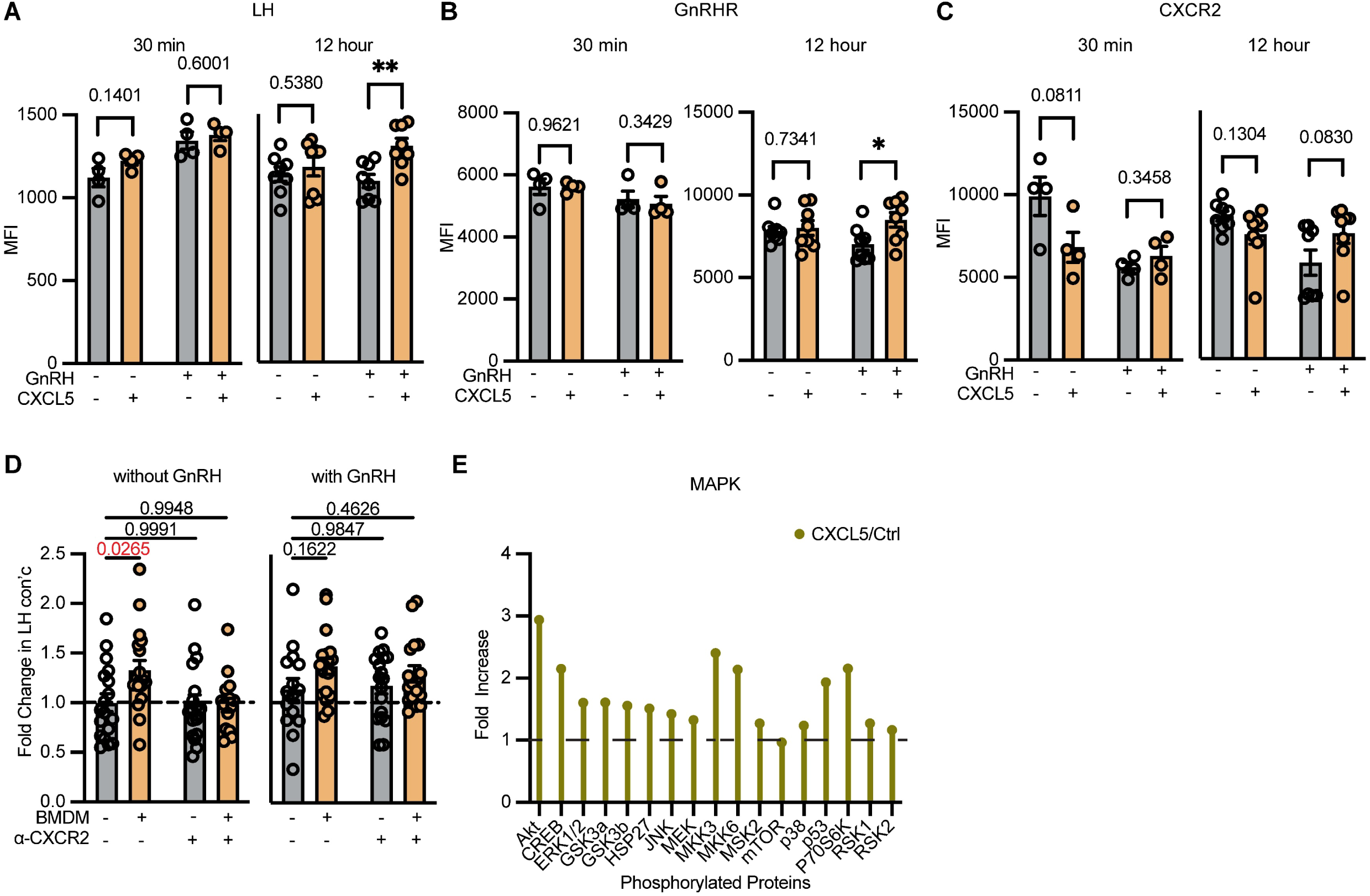
PitMacs modulate LH secretion through CXCL5–CXCR2 activation. (**A**) LH secretion following 12- or 30-hour pre-stimulation with 100 ng/mL CXCL5 and subsequent 30-minute stimulation with 10 nM GnRH is presented as MFI. (**B**-**C**) Corresponding GnRH receptor (GnRHR) and CXCR2 surface expressions were measured by flow cytometry and expressed as MFI. (**D**) LβT2 cells were pre-treated with anti-CXCR2 antibody (1:250 dilution) for 30 minutes prior to 12-hour BMDM-cultured media treatment, with the antibody maintained at a final dilution of 1:500. LβT2 cells were co-cultured with BMDM-conditioned media following the density to volume ratio of 2 gonadotropes per 1 macrophage. LH concentration in culture supernatants was measured by Luminex. (**E**) Phospho-protein array analysis using a mouse MAPK signaling pathway kit. LβT2 cells were treated with 100 ng/mL CXCL5 for 30 minutes without prior serum starvation of cells.

To further understand how gonadotropes dynamically regulate CXCR2 availability in response to CXCL5, we examined CXCR2 surface expression following short-and long-term CXCL5 stimulation. Acute 30-minute CXCL5 stimulation resulted in a trending decrease in CXCR2 expression (**Fig. 5C**), consistent with ligand-induced receptor internalization in chemokine receptors. To confirm that CXCL5 activates gonadotropes through its cognate receptor CXCR2, we pre-incubated LβT2 cells with an anti-CXCR2 blocking antibody prior to 12 hour stimulation with BMDMs. CXCR2 blockade abolished the BMDM-induced increase in LH secretion observed in untreated cells, confirming that secreted molecules from macrophages act specifically through CXCR2 (**Fig. 5D**).

To identify signaling mediators of CXCL5 and IFN-γ activity in LβT2 cells, we profiled phosphorylation events across the MAPK and JAK/STAT pathways (**Fig. 5E, S10C**). Within the MAPK pathway, AKT and ERK were among the proteins with highest levels of phosphorylation, which are the same effectors activated downstream of the GnRHR during gonadotropin stimulation. Collectively, these data demonstrate that CXCL5 signals through CXCR5 and engages the MAPK cascades.

### CXCL5 modulates GnRH-stimulated LH secretion and intracellular LH handling

We next tested the role of MAPK signaling components in CXCL5-induced LH secretion. Given that significant changes in CXCL5-induced LH secretion and GnRHR expression were observed only following 12-hour CXCL5 treatment in GnRH treated conditions, we focused our mechanistic analyses in GnRH-treated samples. Unexpectedly, AKT inhibition increased GnRH-induced LH secretion but had no additive impact on CXCL5-induced LH secretion (**Fig. 6A**). Accompanying this non-additive LH response was a reduction in the GnRHR surface expression upon AKT inhibition of CXCL5 pre-treated cells (**Fig 6B-C**). Interestingly, the transcript level expression of the *Lhb* subunit does not change upon addition of CXCL5 alone, but is significantly reduced upon AKT inhibition, with no statistical effect observed for *Gnrhr* (**Fig. 6D-E**). ERK inhibition had a similar impact on CXCL5-induced LH secretion. ERK inhibition increased GNRH-induced LH secretion with no additive impact on CLC5-induced LH secretion and no changes in GnRHR expression, or *Lhb* and *Gnrhr* transcripts were observed (**Fig. 6G-J).**

**Fig. 6.**
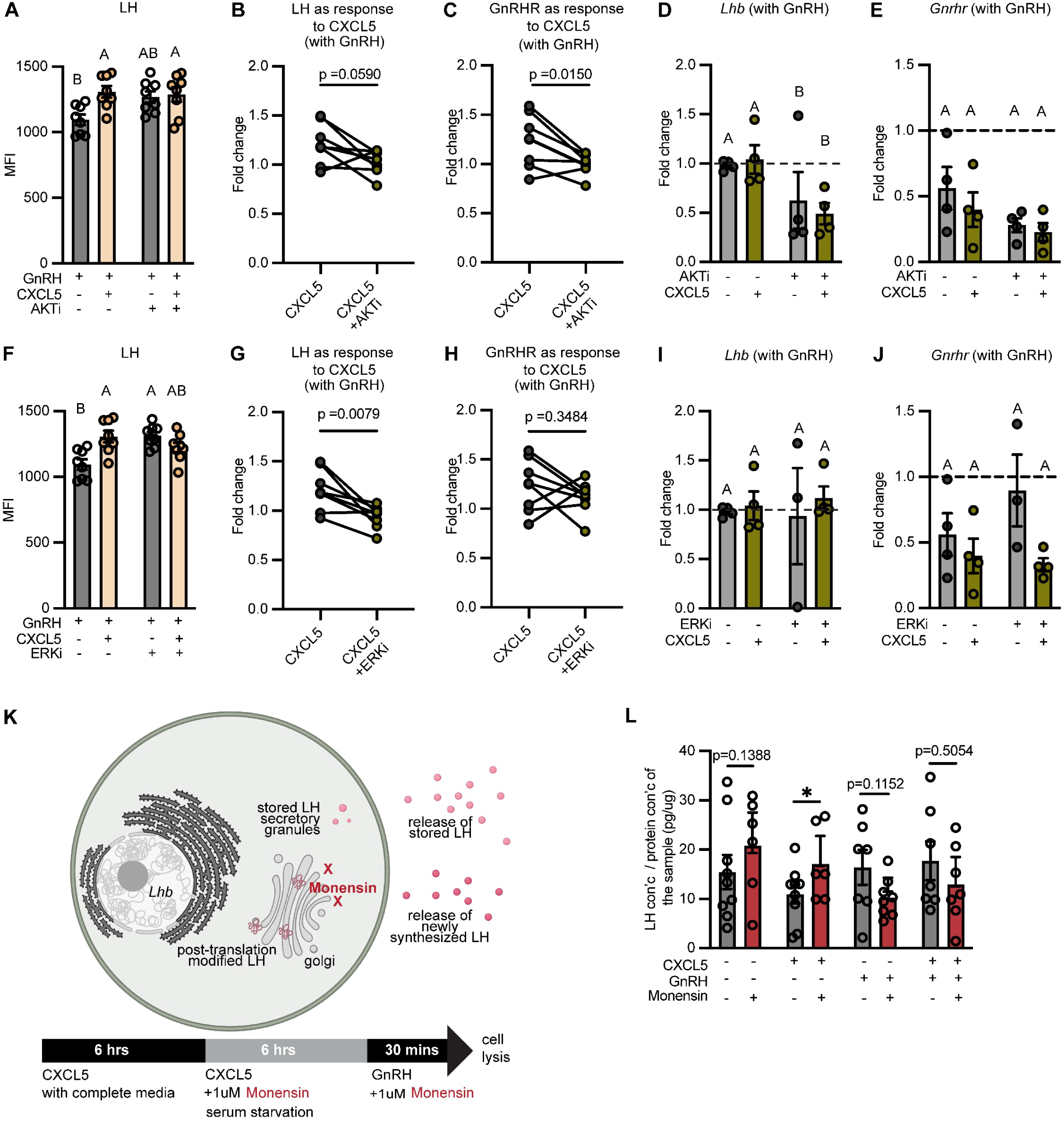
CXCL5 promotes LH secretion through upregulation of synthesis of GnRH-driven release of pre-formed LH. (**A**, **F**) LH secretion following 12-hour pre-stimulation with 100 ng/mL CXCL5 in combination with either (**A**) 20 μM AKT inhibitor or (**F**) 100 nM ERK1/2 inhibitor, and subsequent 30-minute stimulation with 10 nM GnRH, presented as MFI. (**B**, **G**) CXCL5-induced LH secretion expressed as fold change. (**C**, **H**) CXCL5-induced GnRHR surface expression expressed as fold change. qPCR was performed in a 96-well format under similar conditions. (**D**, **I**) *Lhb* and (**E**, **J**) *Gnrhr* expression is shown as fold change from LβT2 cells treated with a combination of 100 ng/mL CXCL5 and either (**D**) 20 μM AKT inhibitor or (**I**) 100 nM ERK1/2 inhibitor. (**K**) Diagram illustrating that cell lysates collected after Monensin treatment contain stored LH and LH that has undergone post-translational modifications and is retained within the cell. LβT2 cells were pre-treated with 100 ng/mL CXCL5 for 12 hours prior to a 30-minute stimulation with 10 nM GnRH. Cells were treated with 1uM of monensin for 6 hours and 30 minutes starting beginning at the 6-hour serum starvation time point. (**L**) Intracellular LH concentration (pg/mL) measured by Luminex assay was normalized to protein concentration of samples determined by BCA assay. Statistical comparisons were performed using Mann-Whitney test or Student’s t-test following outlier removal and normality testing. *p <0.05, **p <0.01, ***p <0.001, ****p <0.0001.

Since reduced *Lhb* transcript levels were discordant with increased LH secretion following CXCL5 stimulation, we asked whether CXCL5 alters the intracellular processing or availability of LH. GnRH stimulates the release of pre-stored LH granules from gonadotropes, whereas newly synthesized LH passes through the Golgi before secretion *(33, 34)*. We therefore used Monensin, an inhibitor of Golgi-dependent protein trafficking, to examine whether CXCL5 alters the intracellular LH pool under basal or GnRH-stimulated conditions *(35)*. LβT2 cells were pre-treated for 12 hours with or without CXCL5, followed by Monensin addition during serum starvation and a 30-minute GnRH stimulation (**Fig. 6K**). Cells were then collected, lysed, and intracellular LH concentration was measured by Luminex.

In the absence of GnRH, Monensin increased intracellular LH in CXCL5-treated cells, whereas the increase in vehicle-treated cells did not reach statistical significance (**Fig. 6L**). This result suggests that CXCL5 alters the Monensin-sensitive intracellular pool of LH under basal conditions, potentially through effects on LH production, processing, trafficking, or turnover. Following GnRH stimulation, Monensin did not significantly alter intracellular LH levels in either vehicle-or CXCL5-treated cells. Thus, although these findings indicate that CXCL5 influences intracellular LH handling, they do not distinguish whether CXCL5 increases de novo LH synthesis, modifies granule trafficking or storage, or alters GnRH-stimulated secretion.

Direct measurements of LH synthesis and secretion kinetics will be required to resolve these possibilities. Notably, in cells treated with both CXCL5 and GnRH, intracellular LH levels were unchanged by Monensin, suggesting that CXCL5 pre-treatment augments LH synthesis while GnRH simultaneously promotes the release of stored LH, resulting in a net balance between production and secretion.

## DISCUSSION

In this study, we demonstrate that PitMacs are transcriptionally and functionally distinct from other tissue-resident macrophages and serve a pituitary-specific role in modulating female hormone secretion through cytokine-mediated signaling.

The pituitary gland’s anatomical position at the base of the brain is frequently underappreciated from an immunological standpoint. Unlike the central nervous system parenchyma, the pituitary resides outside the blood-brain barrier and is therefore accessible to peripheral immune surveillance, rendering it a unique neuroimmune regulatory microenvironment *(36-38)*. Here, we demonstrate that PitMacs constitute the dominant immune cell population within this tissue and are transcriptomically distinct from microglia, the canonical brain-resident macrophage whose identity, heterogeneity, and function have been extensively characterized *(39, 40)*. Differential Gene Set Enrichment Analysis (DGSEA) of pituitary macrophage subsets revealed transcriptional associations with hormone production, interferon (IFN) response, and heat shock responses. Given that hormone secretion is a defining function of the pituitary gland, we investigated the contribution of PitMacs to endocrine hormone modulation. Within our integrated scRNA-seq dataset, *Gh* and *Prl* emerged among the top differentially expressed genes in PitMacs, and our RiboTag experiment further confirmed their active translation. Although our *in vivo* and *in vitro* macrophage depletion experiments revealed trending decreases in GH and PRL secretion, the inconsistent statistical significance suggests that macrophage-derived hormone production occurs at quantities substantially lower than those generated by the canonical endocrine cell populations, somatotropes and lactotropes. Nevertheless, even modest macrophage-derived hormone production may contribute meaningfully to neuroendocrine crosstalk. Both GH and PRL belong to the class I cytokine receptor superfamily and therefore possess cytokine-like functions *(41)*. PRL has previously been shown to be secreted by immune cells, including macrophages and T cells, where it promotes cytokine secretion and activates the STAT1 and MAPK pathways *(42-46)*. Furthermore, PRL can induce nitric oxide production and tumor necrosis factor-α (TNF-α) secretion in murine peritoneal macrophages and plays an autocrine role in T cell growth and activation *(47, 48)*. GH, in turn, has been shown to enhance Th1 cytokine activity, including production of IFN-γ *(49)*.

To further support the potential for crosstalk between PitMacs and endocrine cells, we demonstrated that PitMacs actively translate hormone receptor transcripts, specifically *Gnrhr* and *Ghrhr*. Within the pituitary, GnRHR is canonically expressed in gonadotropes, enabling responsiveness to pulsatile hypothalamic GnRH signals, but is also present in extra-pituitary tissues including the ovary, breast, lymphocytes, and mast cells *(50-52)*. GHRHR, while primarily expressed in somatotropes to regulate growth hormone secretion, is similarly expressed in a range of peripheral cell types, including non-malignant and malignant cells and immune cells *(53, 54)*. This observation underscores the importance of investigating the immunological role of PitMacs in the context of hormone signaling.

The pituitary-specific function of macrophages is further supported by depletion experiments conducted both *in vitro* and *in vivo*. Despite comprising only approximately 5% of the total pituitary cell population, macrophage depletion significantly reduced hormone secretion. Notably, in our AAV-based *in vivo* model, depletion of only 23.48% of the pituitary macrophage pool was sufficient to produce significant decreases in serum LH and FSH in mice with an intact hypothalamic-pituitary-gonadal (HPG) axis. This disproportionate impact of modest macrophage loss may reflect the highly organized and interdependent nature of the pituitary cellular network, in which endocrine and non-endocrine pituitary cells such as folliculostellate cells and vascular cells engage in robust cell–cell interactions. Disruption of even a minor cellular fraction may therefore be sufficient to impair coordinated hormone secretion *(55-58)*.

We further propose a mechanism by which PitMacs promote gonadotropin secretion, centered on cytokine-mediated interactions with gonadotropes via CXCL5. CXCL5 signals through CXCR2 and engages multiple downstream pathways, with the MAPK cascade serving as a shared signaling node in gonadotropes *(59-61)*. We demonstrate that gonadotropes pre-treated with AKT or ERK inhibitors exhibit LH secretion levels comparable to those treated with CXCL5 alone, suggesting pathway convergence rather than additive signaling. One mechanistic explanation is that CXCL5 stimulation upregulates RAF activity upstream of ERK, while AKT inhibition concurrently relieves AKT-mediated suppression of RAF, such that CXCL5 alone or AKT inhibition combined with GnRH stimulation are each sufficient to drive LH secretion through the RAF–ERK axis *(61-63)*.

Furthermore, our findings demonstrate that CXCL5-induced LH secretion is time-dependent in its relationship with GnRH. At 30 minutes, LH release proceeds in a GnRH-independent manner, whereas by 12 hours it becomes GnRH-dependent, suggesting that sustained CXCL5 signaling progressively engages GnRH-coupled pathways. These temporal changes in LH secretion were accompanied by concurrent alterations in GnRHR surface expression, recapitulating the well-described plasticity of pituitary gonadotropes with respect to GnRHR expression and receptor density across reproductive cycles *(64, 65)*. Additionally, acute CXCL5 exposure drove CXCR2 internalization and transient receptor desensitization, whereas sustained CXCL5 stimulation permitted receptor recovery and established a primed cellular state in which GnRH could actively upregulate CXCR2. Collectively, these findings provide evidence for signaling convergence between the CXCL5-CXCR2 and GnRH-GnRHR axes in gonadotropes.

While we demonstrate that both CXCL5 and IFNγ can be produced by PitMacs, our data also indicate that pituitary T cells are significant contributors to these cytokines, implicating a broader immune cell network in the regulation of gonadotropin secretion. Macrophage–T cell crosstalk may amplify this signaling cascade, as macrophage-derived IL-2 can promote T cell production of IFNγ. Additionally, CXCL5 may originate from pituitary-resident neutrophils, further expanding the repertoire of potential immune sources. The small amounts of Prl and GH translated by PitMacs may also act in a paracrine manner to modulate neighboring immune or endocrine cells within the pituitary microenvironment. To more precisely delineate the relative contributions of distinct immune cell populations to gonadotrope function, future studies employing targeted depletion of specific immune subsets followed by assessment of gonadotropin secretory responses will be essential.

Our RiboTag data further revealed significant active translation of *Cga* and *Pomc* transcripts in PitMacs. *Cga* encodes the common α-subunit shared by TSH, LH, FSH, and hCG, whereas *Pomc* is the precursor of ACTH, α-MSH, and β-endorphin. The active translation of these hormone subunits and precursors within PitMacs suggests a previously unappreciated complexity in their interactions with hormone-producing endocrine cells. In addition, although our primary cytokine analyses centered on IFN-γ, we detected *Ifnar1* expression in PitMacs, suggesting a contribution of type I IFN signaling within this compartment. Notably, our flow cytometry data indicate that PitMacs are the predominant producers of IFN-β among pituitary immune cells, surpassing T cells. This finding was not captured by our cytokine multiplex panels, which do not include IFN-β. Taken together with our transcriptomic findings and those of Lehtonen et al., who described an interferon-associated macrophage subset in the pituitary, type I IFN signaling between PitMacs and endocrine cells represents an important and underexplored axis of neuroendocrine-immune regulation.

In this study, we demonstrate that PitMacs are transcriptionally and functionally distinct from other tissue-resident macrophages, uniquely shaped by the endocrine microenvironment of the pituitary gland. Residing outside the blood-brain barrier, the pituitary is accessible to immune surveillance, positioning PitMacs as active participants in hormonal output through cytokine-mediated signaling and immune cell crosstalk with direct consequences for gonadotropin secretion and HPG axis function. These findings provide new insights into the immune regulation of reproductive physiology and establish a foundation for understanding how pituitary inflammation contributes to hormonal disorders, bridging immunology and pituitary biology.

## MATERIALS AND METHODS

### Mice

C57BL/6J mice were obtained directly from The Jackson Laboratory (Stock No. 000664) and used for experiments as described. A separate breeding colony of C57BL/6J mice was also maintained in the McGaugh Hall Vivarium at the University of California, Irvine. LysMCre (B6.129P2-Lyz2tm1(cre)Ifo/J; The Jackson Laboratory, Stock No. 004781) and RiboTag (B6J.129(Cg)-Rpl22tm1.1Psam/SjJ; The Jackson Laboratory, Stock No. 029977) mice were each maintained as independent breeding colonies in the same vivarium. LysMCre and RiboTag mice were intercrossed at two females to one male per breeding cage to generate LysMCre × RiboTag offspring for use in RiboTag experiments. All mice were housed under a 12-hour light/dark cycle with ad libitum access to standard rodent chow and water. All experimental procedures were performed in accordance with the National Institutes of Health Guide for the Care and Use of Laboratory Animals and approved by the University of California, Irvine Institutional Animal Care and Use Committee (IACUC Protocol No. AUP-24-068).

### Single-Cell Secretome Profiling by Isoplexis

Pituitaries were harvested from two 10-week-old male and two 10-week-old female C57BL/6J mice. Pituitaries from male and female mice were independently dissociated into single-cell suspensions (see in vitro macrophage depletion). Dissociated cells were stimulated with 10 μg/mL lipopolysaccharide (LPS) for 24 hours. Stimulated cells were loaded onto the Isoplexis platform and incubated overnight in the Isoplexis chamber to capture single cell secretome profiles according to the manufacturer’s instructions.

### *In Vivo* Macrophage Depletion by AAV-Caspase

To achieve targeted in vivo depletion of pituitary macrophages, 1 × 10¹² viral genome particles of an adeno-associated virus (AAV) encoding an auto-activating caspase driven by a Cre-dependent, GH promoter-controlled construct were administered via retro-orbital injection to 10-week-old female LysMCre mice. An AAV encoding GFP in place of caspase was used as the control. Estrous cyclicity was monitored daily beginning 1 week prior to injection and continuing for 3 weeks post-injection (see Estrous Cycle Assessment). GnRH challenge was performed immediately prior to AAV injection and weekly thereafter for 3 weeks to collect serum for measurement of circulating LH and FSH levels (see GnRH Challenge and Serum Collection). Pituitaries were collected at experimental endpoint for flow cytometric confirmation of macrophage depletion.

### GnRH Challenge

Each mouse received an intraperitoneal injection of GnRH (150 ng/kg) dissolved in sterile PBS. Approximately 50 μL of blood was collected via the retro-orbital sinus at baseline (pre-GnRH) and at 15 minutes post-injection. All injections and blood collections were performed under isoflurane inhalant anesthesia

### Serum Collection

Blood samples were collected weekly for three weeks following AAV injection, with both pre-and post-GnRH samples obtained at each time point. All samples, except for the post-GnRH week 3 collection, were obtained via the retro-orbital sinus. The week 3 post-GnRH sample was collected from trunk blood following euthanasia. All blood samples were allowed to clot at room temperature for 1 hour, centrifuged at 2,000 × g for 10 minutes, and serum was collected and stored at −20°C until assayed.

### In Vitro Macrophage Depletion

Pituitaries from 10-week-old female C57BL/6J mice were dissociated into single-cell suspension by mechanically passed through 100uM cell filter followed by washes of complete DMEM media. Tissue was then treated with DNAse I (StemCell Technologies, Cat. No. 07900) for 10 minutes at 37°C. Tissue was then washed and resuspended in FACS buffer. Macrophages were depleted from pooled pituitary single-cell suspensions using the EasySep™ Mouse F4/80 Positive Selection Kit (STEMCELL Technologies, Cat. No. 100-0659) according to the manufacturer’s instructions. Both dispersed and macrophage-depleted pituitary cells were plated and cultured in complete DMEM supplemented with 10% FBS at 37°C in a humidified 5% CO₂ incubator for 12 hours prior to any treatment. This protocol was used for all primary pituitary rescue experiments described herein.

### BMDM Rescue and anti-CXCR2 Treatment

Following 12 hours of culture in complete media post-plating, LβT2 cells were pre-treated with anti-CXCR2 antibody (ThermoFisher Scientific, Cat. No. PA5-102662) at a 1:250 dilution for 30 minutes at 37°C. Cells were subsequently treated with either BMDM-conditioned media or complete media for 6 hours, bringing the final anti-CXCR2 antibody dilution to 1:500. BMDM-conditioned media was collected from 12-hour BMDM in vitro cultures. Following the 6-hour treatment, media was replaced with serum-free DMEM containing the same treatments. After an additional 6 hours of serum starvation, media was replaced with serum-free DMEM with or without 10 nM GnRH for 30 minutes to perform *in vitro* GnRH stimulation. Supernatants were collected and stored at −20°C until used for Luminex hormone measurements.

### Phosphoprotein Array

LβT2 gonadotrope cells were stimulated with either 100 pg/mL IFN-γ or 100 ng/mL CXCL5 for 30 minutes. LβT2 cells stimulated with 10 nM GnRH and BMDMs stimulated with 100 pg/mL IFN-γ served as controls. Following stimulation, cells were lysed in RIPA buffer supplemented with protease and phosphatase inhibitor cocktails. Total protein concentrations were determined by BCA assay. Antibody-printed membranes from the JAK/STAT and MAPK Phospho Antibody Array kits (RayBiotech, Cat. No. AAM-JAKSTAT-1-4, Cat. No. AAH-MAPK-1-4) were incubated with normalized cell lysates overnight at 4°C, followed by sequential incubation with biotinylated detection antibodies and HRP-conjugated streptavidin as recommended by the manufacturer. Chemiluminescent signal was detected using a Bio-Rad ChemiDoc Imaging System, and densitometry was quantified using ImageJ.

### MAPK Pathway Inhibitor Treatment

LβT2 cells were cultured in complete media for 12 hours prior to treatment. Cells were then treated with a combination of 100 nM ERK1/2 inhibitor, 20 μM AKT inhibitor, and 100 ng/mL CXCL5 for 6 hours in complete media, followed by an additional 6 hours in serum-free media. Cells were subsequently stimulated with 10 nM GnRH for 30 minutes. Supernatants were collected and stored at −20°C for future Luminex LH analysis. For flow cytometric analysis of surface GnRHR and CXCR2 expression, LBT2 cells were seeded in 24-well plates at 4.0 × 10⁵ cells/mL in 500 μL per well. For RNA extraction and qPCR quantification of Lhb expression, LBT2 cells were seeded in 96-well plates at 1.3 × 10⁵ cells/mL in 50 μL per well and processed as described in the RNA Extraction and qPCR section.

### BMDM and Pituitary Macrophage Rescue

Following 12 hours of culture in complete media post-plating, macrophage-depleted pituitary cells were treated with conditioned media from, or co-cultured directly with, either BMDMs or pituitary-resident macrophages. Co-culture cell numbers were established at a ratio of 2 gonadotropes per 1 macrophage. At 6 hours post-treatment, media was replaced with serum-free DMEM containing equivalent treatment conditions. After an additional 6 hours of serum starvation, media was replaced with serum-free DMEM with or without 10 nM GnRH to perform in vitro GnRH stimulation. Supernatants were collected and stored at −20°C until used for Luminex LH measurements. BMDM and pituitary macrophage conditioned media were harvested from 12-hour *in vitro* cultures. Experiments were conducted in both 96-and 384-well plate formats. LH concentrations across all wells were normalized to the mean LH concentration of non-GnRH-stimulated dispersed pituitary wells within each experiment to calculate fold change.

### LBT2 Dose and Time Response

LβT2 cells were seeded in 96-well plates at a density of 1.28 x 10^6^ cells/mL in 50 μL per well. Cells were treated with CXCL5 (0.1, 1, 10, or 100 ng/mL) or IFN-γ (1, 10, 100, or 1,000 pg/mL) for 12 hours. Conditioned-media collected at each time point and stored at −20°C until used for Luminex LH measurements.

### RiboTag-seq

Pituitaries from 9-to 10-week-old heterozygous LysMCre × RiboTag female mice were collected exclusively during diestrus, immediately flash-frozen in liquid nitrogen, and stored at −80°C until processing. Pituitaries were pooled, homogenized, and lysed, and HA-tagged ribosome-associated RNA complexes were immunoprecipitated using Pierce Anti-HA Magnetic Beads (Thermo Fisher, Cat. No. 88836) according to the manufacturer’s instructions. RNA was extracted from both total input and immunoprecipitated (IP) fractions using the RNeasy Micro Kit (Qiagen, Cat. No. 175013887). Libraries were prepared using a stranded rRNA depletion library preparation protocol and sequenced on the Illumina NovaSeq X Plus platform.

### RiboTag-seq Data Analysis

Raw sequencing data from male and female input and IP samples were assessed for quality using FastQC and adapter-trimmed using fastp. Trimmed reads were aligned to the mouse reference genome (GRCm38/mm10) using STAR with a genome index generated from Ensembl annotation. Read counts were quantified using DESeq2 and exported as a count matrix for downstream differential expression analysis performed in RStudio.

### Cytokine and Hormone Multiplex Immunoassay

A panel of 27 cytokines was quantified using the Milliplex MAP Mouse Cytokine/Chemokine Magnetic Bead Panel (Millipore Sigma, Cat. No. MCYTOMAG-70K) according to the manufacturer’s instructions. Analyte concentrations were measured on the INTELLIFLEX instrument (Luminex) and analyzed using BELYSA software (v1.2). Similarly, hormones were quantified using the Milliplex MAP Mouse Pituitary Panel (Millipore Sigma, Cat. No. MPTMAG-49K, to measure LH, FSH, Prl, and GH.

### Quantification and Statistical Analysis

All data are presented as mean ± SEM. Statistical analyses were performed using JMP Student Edition 19. Datasets were assessed for normality prior to group comparisons, and Box-Cox transformation was applied to non-normally distributed data to satisfy the assumptions of parametric testing. Outliers were identified and removed using the ROUT method (Q = 1%) in GraphPad Prism 9. Group differences were evaluated by one-way or two-way ANOVA, as appropriate for each experimental design, followed by post-hoc Tukey’s multiple comparisons test or Dunnett’s test for comparisons against a single reference group. All graphs were generated in GraphPad Prism 9. A p-value of ≤ 0.05 was considered statistically significant. In figures where compact letter display is used, groups not sharing a common letter differ significantly from one another. Exact sample sizes, statistical tests used, and p-values for each experiment are reported in the corresponding figure legends.

### Estrous Cycle Assessment

Mouse vaginal cytology was performed daily to monitor estrous cyclicity beginning 1 week prior to AAV injection and continuing for 3 weeks post-injection. Estrous cycle stages were classified based on the predominant cell population observed: proestrus, by nucleated epithelial cells; estrus, by cornified anucleated epithelial cells; metestrus, by a mixture of cornified epithelial cells, nucleated epithelial cells, and leukocytes; and diestrus, by the predominance of leukocytes.

### RNA extraction and qPCR

RNA was extracted from LβT2 cells treated with CXCL5 and inhibitors (see MAPK Pathway Inhibitor Treatment) using the RNAeasy 96 kit (Qiagen, Cat. No. 74181). cDNA was synthesized using the iScript Kit (BioRad, Cat. No. 1708841). RT-qPCR was performed using the iQ SYBR Green Supermix Kit (BioRad, Cat. No. 1708882). Relative gene expressions of *Lhb* and *Gnrhr* were calculated using the 2-ΔΔCq method and *Gapdh* as the reference gene.

### Intracellular LH Measurement Post-Monensin Treatment

LβT2 cells were cultured in 24-well plates at a density of 0.8 x 10^5^ cells/mL in complete media for 12 hours prior to treatment. Cells were then treated with or without 100 ng/mL CXCL5 for 6 hours in complete media, followed by an additional 6 hours in serum-free media. Subsequently, cells were stimulated with 10 nM GnRH for 30 minutes. 1uM of Monensin was added during the 6 hour serum starvation and 30-minute GnRH stimulation. After treatment, cell lysates were collected to measure total protein concentration using BCA assay and LH concentration using the Luminex LH assay.

### ScRNASeq Integration

Two mouse single-cell RNA sequencing (scRNA-seq) datasets were analyzed including the publicly available Tabula Muris atlas and an in-house pituitary gland dataset generated by our laboratory investigating immune cell dynamics in control and letrozole-induced polycystic ovary syndrome (PCOS) conditions. All analyses were conducted in R using the Seurat framework (v4). Raw feature names in the Tabula Muris dataset, originally encoded as Ensembl gene identifiers, were converted to MGI gene symbols using the biomaRt package by querying the Ensembl BioMart database (mmusculus_gene_ensembl). Both datasets were restricted to cells from 3-month-old mice. The Tabula Muris dataset was further subset to 22 tissues of interest including large intestine, aorta, heart, brain, diaphragm, pancreas, brown adipose tissue, tongue, mammary gland, lung, skin, spleen, liver, kidney, subcutaneous adipose tissue, thymus, bone marrow, trachea, gonadal fat pad, limb muscle, bladder lumen, and mesenteric fat pad. Our laboratory’s pituitary dataset was restricted to control-treated animals and immune cell populations. Cluster labels were manually assigned based on canonical marker gene expression lists from PanglaoDB. Tissue-resident macrophages within the Tabula Muris dataset were further annotated with tissue-specific identities (e.g., lung macrophage, Kupffer cell, heart macrophage, bone marrow macrophage) by cross-referencing cell type labels with corresponding tissue metadata.

Subsetted immune cell populations from both datasets were merged into a single Seurat object, with dataset of origin, tissue identity, and disease status recorded in cell metadata. The merged object was log-normalized, and the top 2,000 highly variable features were identified with the variance-stabilizing transformation method. All genes were subsequently scaled, and principal component analysis (PCA) was performed on the top variable features retaining 22 principal components. Batch effects attributable to dataset of origin were mitigated using Harmony integration, with algorithmic convergence confirmed visually. A shared nearest-neighbor graph was then constructed from the Harmony-corrected embeddings across 22 dimensions,and unsupervised clustering was performed with the Louvain algorithm at a resolution of 0.5.

Broad marker identification across all macrophage and immune cell populations was carried out using restricted to positively expressed genes. Differentially expressed genes (DEGs) were filtered using a log2 fold-change threshold greater than 1 and a nominal p-value less than 0.05, and full DEG lists were exported as CSV files. Heatmaps of the top 5 DEGs per cluster were produced using DoHeatmap() with custom diverging color gradients. Targeted heatmaps were additionally generated by intersecting cluster-level DEGs with gene sets representing macrophage functional programs, including phagocytosis, activation, and migration from the Gene Ontology Resource.

## Funding

National Institutes of Health Training Grant T32 AI 177324

*Eunice Kennedy Shriver* National Institute of Child Health and Human Development R00HD098330

Graduate Assistance in Areas of National Need (GAANN) training grant (P200A210024)

## Author Contributions

Conceptualization: ZDM, DN

Methodology: ZDM, SC, EVV, DSK, DN, KT

Investigation: ZDM, JH, LZ, AZ, GDR, KW, KP, NU, JAA, KT

Visualization: ZDM, JH, GDR, DN

Supervision: ZDM, DN

Writing—original draft: ZDM, DN

Writing—review & editing: ZDM, JH, LZ, AZ, GDR, KW, KP, NU, JAA, KT, SC, EVV, DSK, DN

