## Supplemental Figures for "Endocrine-Adapted Pituitary Macrophages Regulate Gonadotropin Secretion through CXCL5-CXCR2-MAPK Signaling"

**Supplemental Figures S1-S10**
**Supplemental Methods**

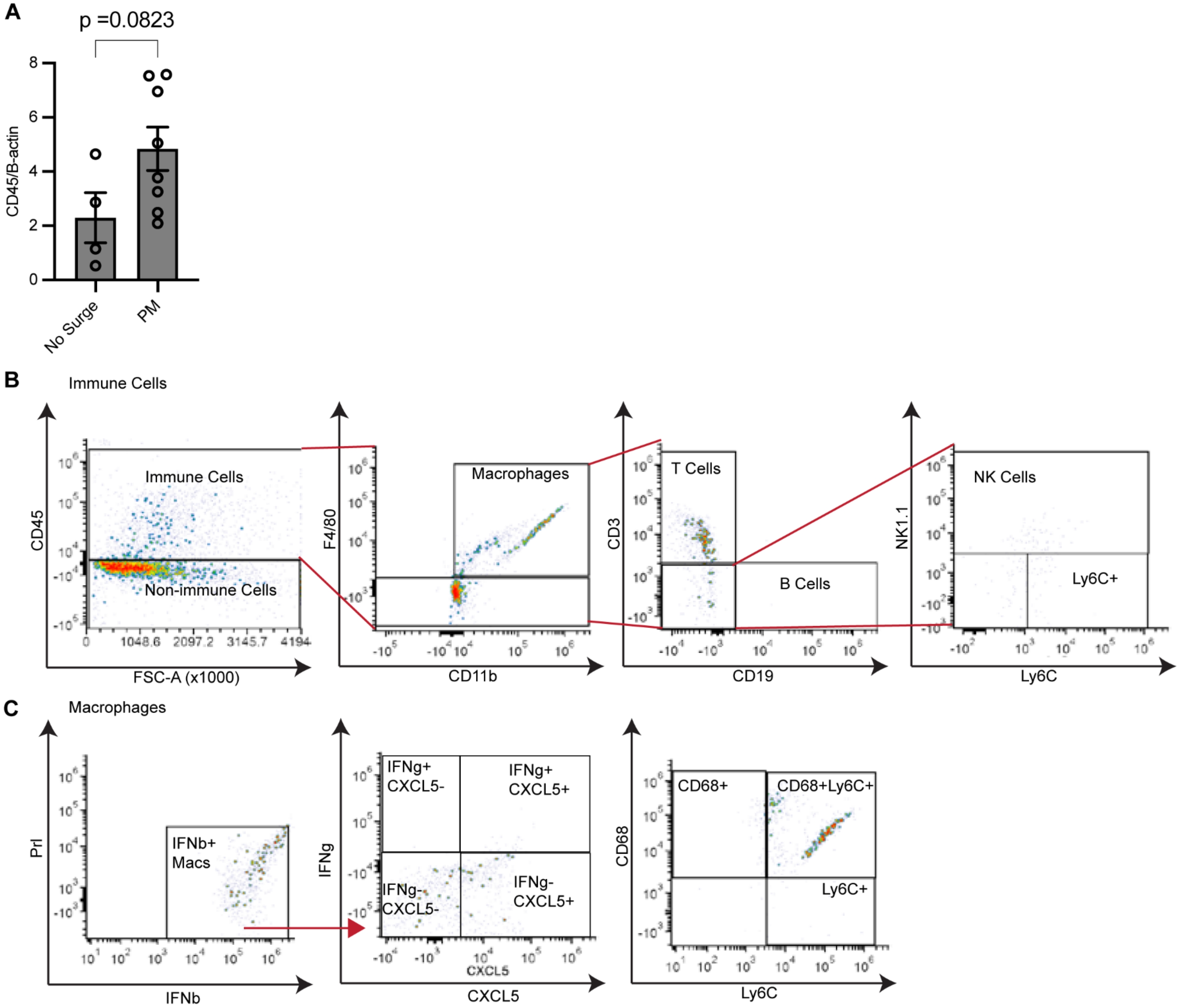


**Supplementary Figure 1.** (**A**) Quantification of western blot showing CD45 expression from OVX+E2-treated mice during LH surge, further divided by mice that did and did not exhibit induced LH surge. CD45 expression normalized to B-actin, presented as fold expression. (**B**) Flow cytometry gating strategy for immune cell populations isolated from primary mouse pituitaries. (**C**) Macrophages (F4/80+CD11b+) were further gated for intracellular cytokines, IFNb, IFNg, and CXCL5, as well as surface markers and CD68 and Ly6C.

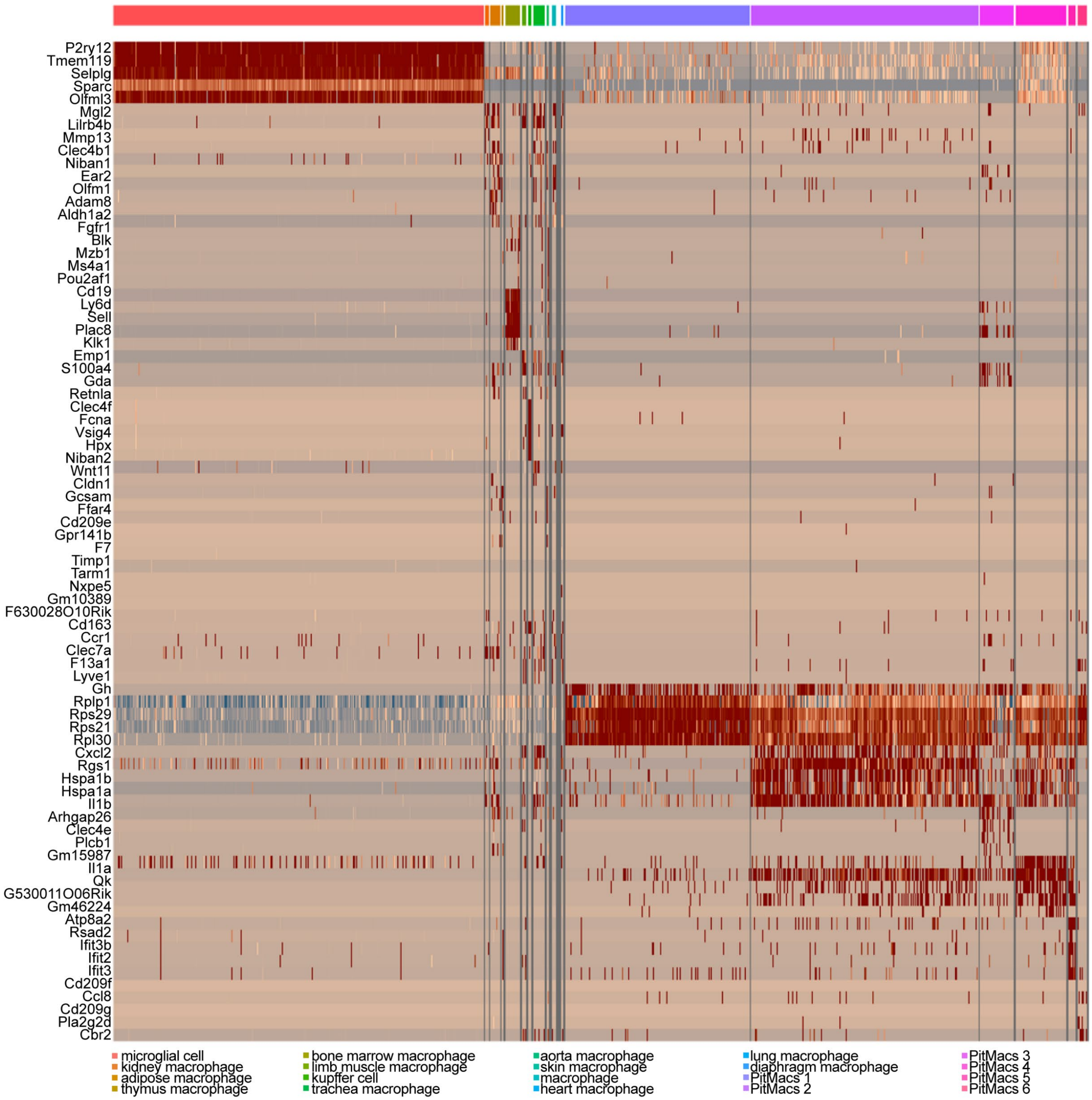


**Supplementary Figure 2.** Heatmap displaying the top differentially expressed genes (DEGs) in macrophages from distinct tissue depots in the integrated Tabula Muris and pituitary scRNA-seq datasets.

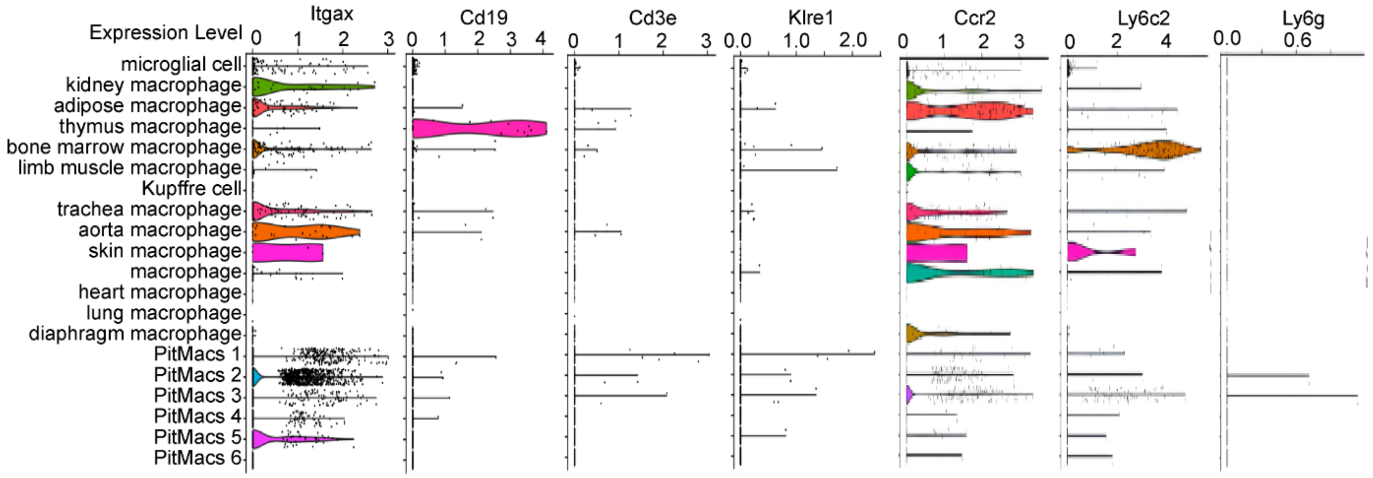


**Supplementary Figure 3.** Violin plots displaying the expression levels of immune canonical markers *Itgax, Cd19, Cd3e, Klre1, Ccr2, Ly6c2,* and *Ly6g* in macrophages from distinct tissue depots.

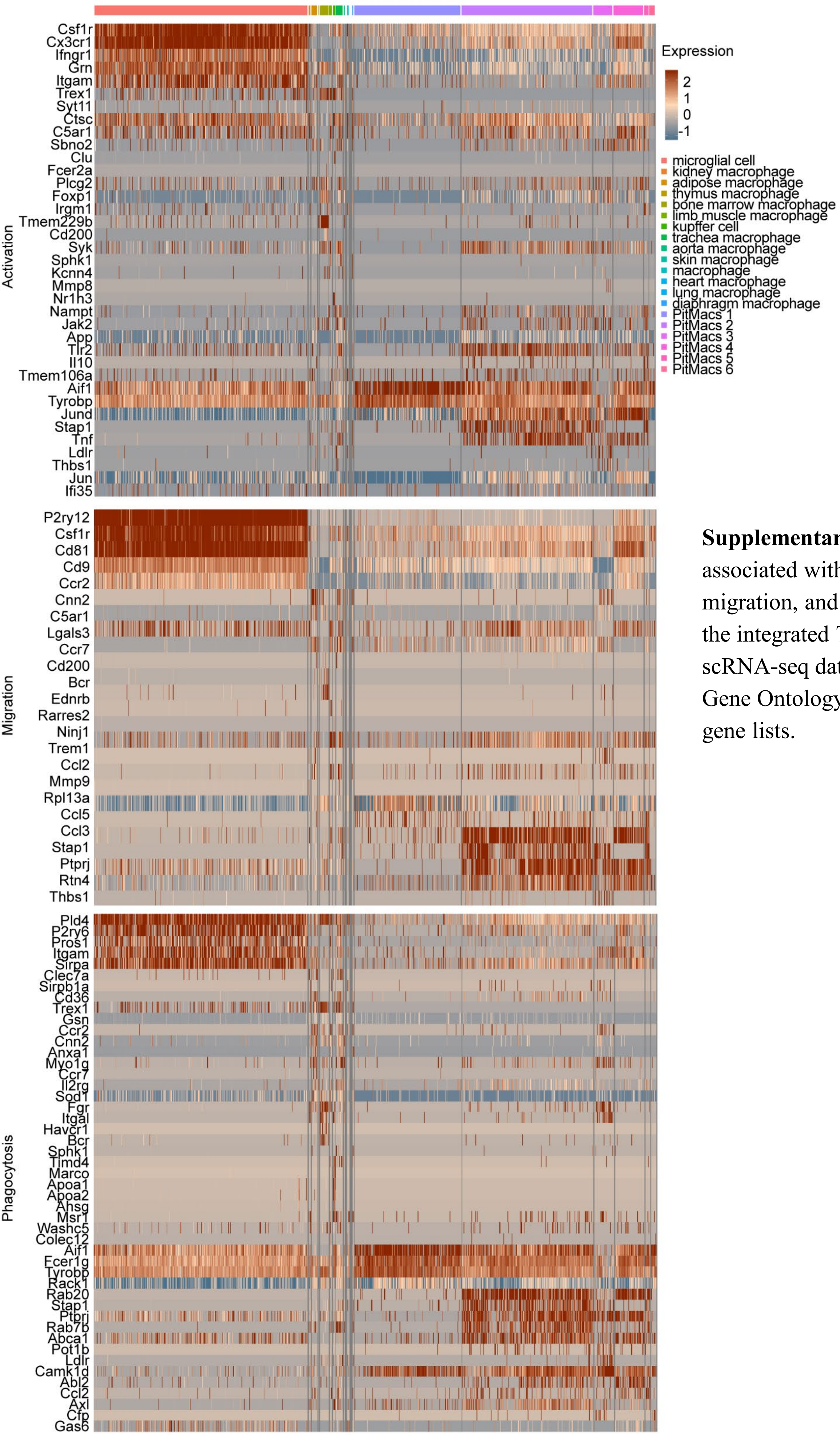


**Supplementary Figure 4.** Genes associated with macrophage activation, migration, and phagocytosis. DEGs from the integrated Tabula Muris and pituitary scRNA-seq datasets were filtered against Gene Ontology (GO) macrophage function gene lists.

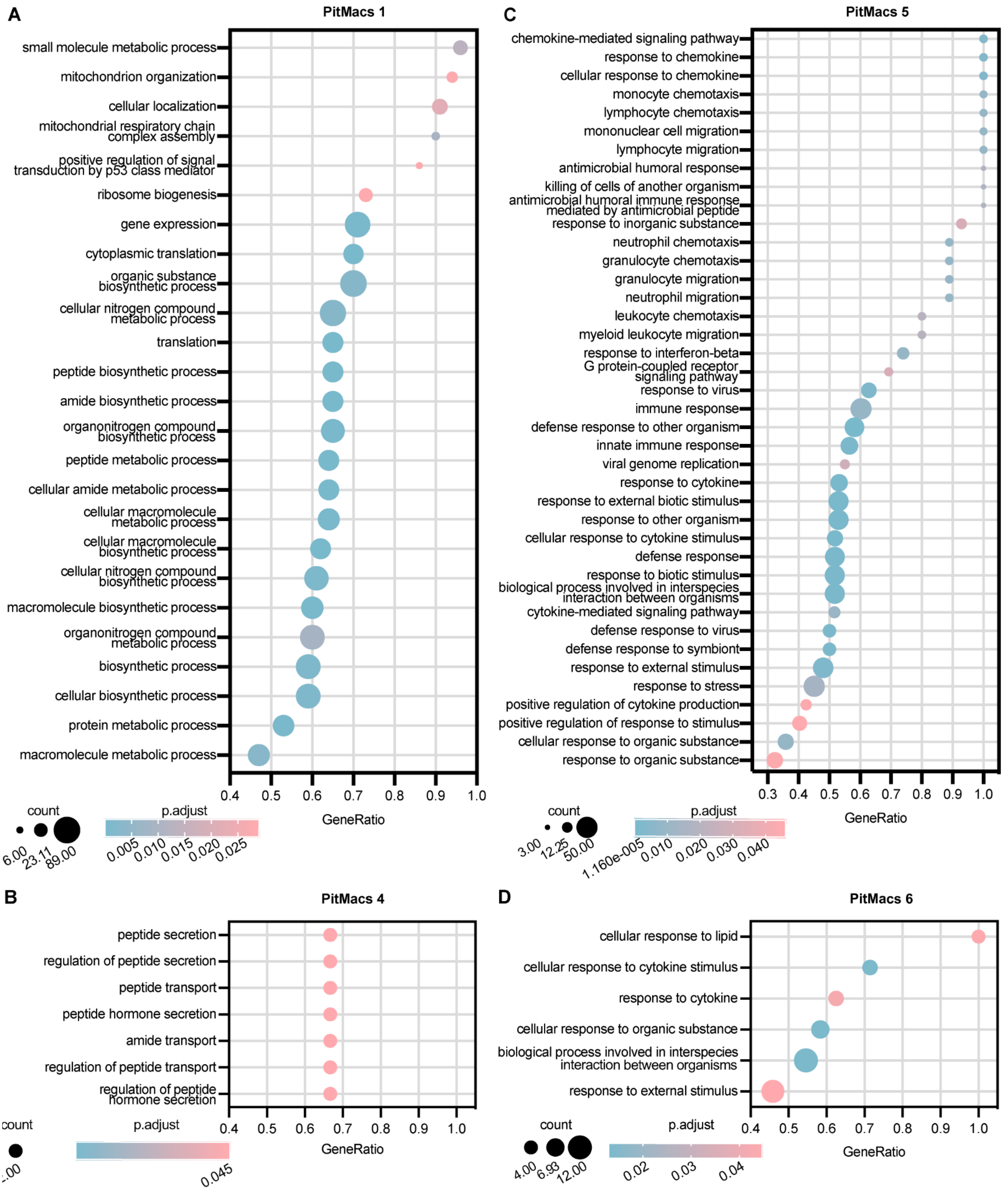


**Supplementary Figure 5.** Differential Gene Set Enrichment Analysis (DGSEA) was performed using the differentially expressed genes (DEGs) identified within each PitMac cluster. Dot plots illustrate the Gene Ontology (GO) terms significantly enriched in each cluster, with enrichment significance quantified by p-adjusted values and gene counts. (**A**) PitMac 1; (**B**) PitMac 4; (**C**) PitMac 5; (**D**) PitMac 6. No significantly enriched GO terms were identified for the remaining PitMac clusters.

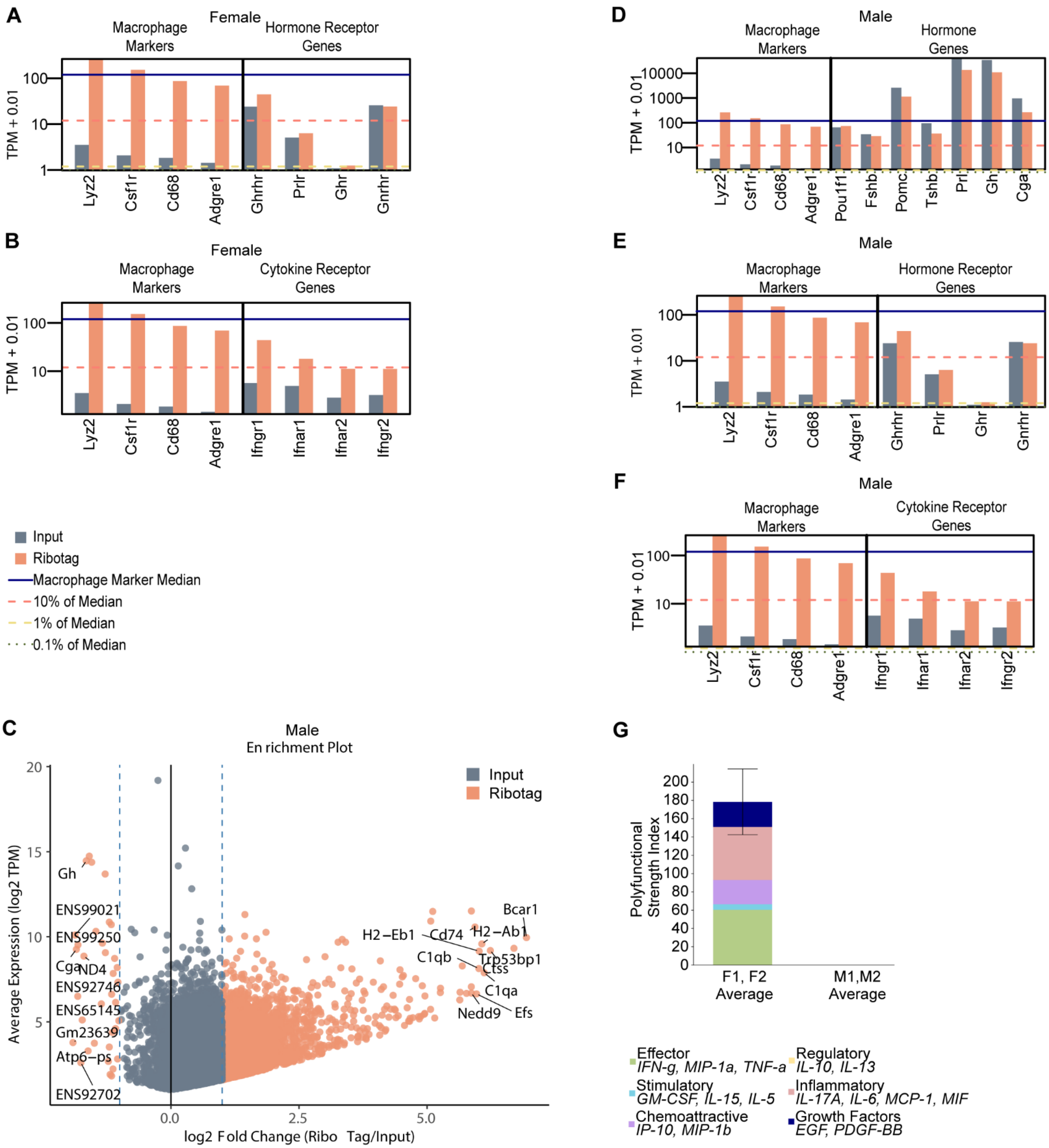


**Supplementary Figure 6.** Expression levels of canonical macrophage markers (Lyz2, Csf1r, Cd68, and Adgre1) are shown alongside genes of interest encoding hormone receptors (**A, E**) hormones (**D**), and cytokine receptors (**B, F**). Analysis was done separately for fractions from female **(A-B)** and male mice (**D-F**). Expression values are presented as Transcripts Per Million (TPM), with the percent median TPM of macrophage markers used as a baseline reference (**C**) Enrichment plot showing genes highly enriched in the immunoprecipitated (RiboTag) fraction relative to total input in male mouse samples. (**G**) PitMacs isolated from 10-week-old female and male mice were pre-treated with 10 µg/mL LPS for 24 hours, followed by overnight incubation in LPS-free media to characterize single-cell secretome profiles. Stacked bars represent different cytokine categories and is presented polyfunctional strength indices from n=2 female (F1,F2) and n=3 male (M1, M2) PitMacs.

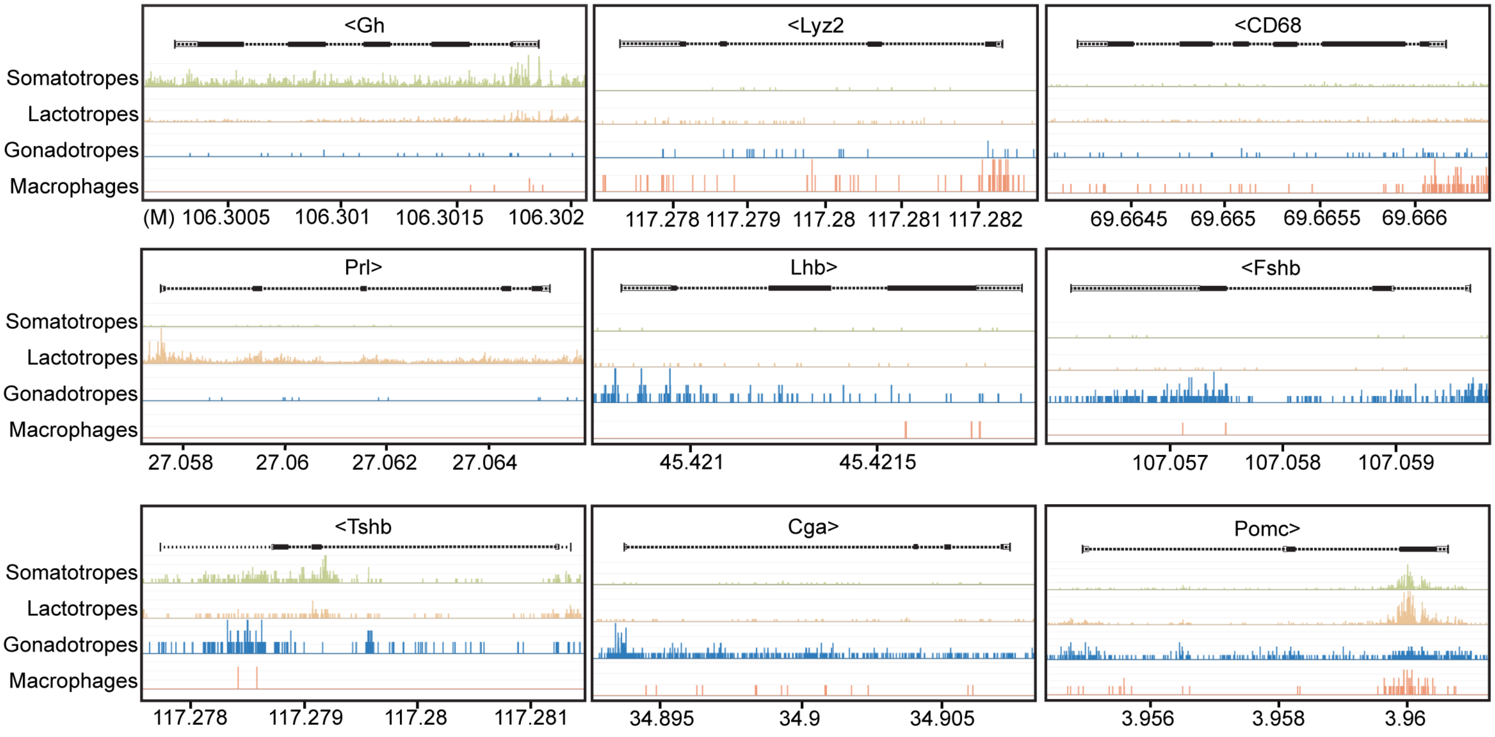


**Supplementary Figure 7.** ATAC-seq data from Wallis et al. demonstrating open chromatin at the *Gh* locus in PitMacs (macrophages). Gene markers for pituitary endocrine cell types are shown.

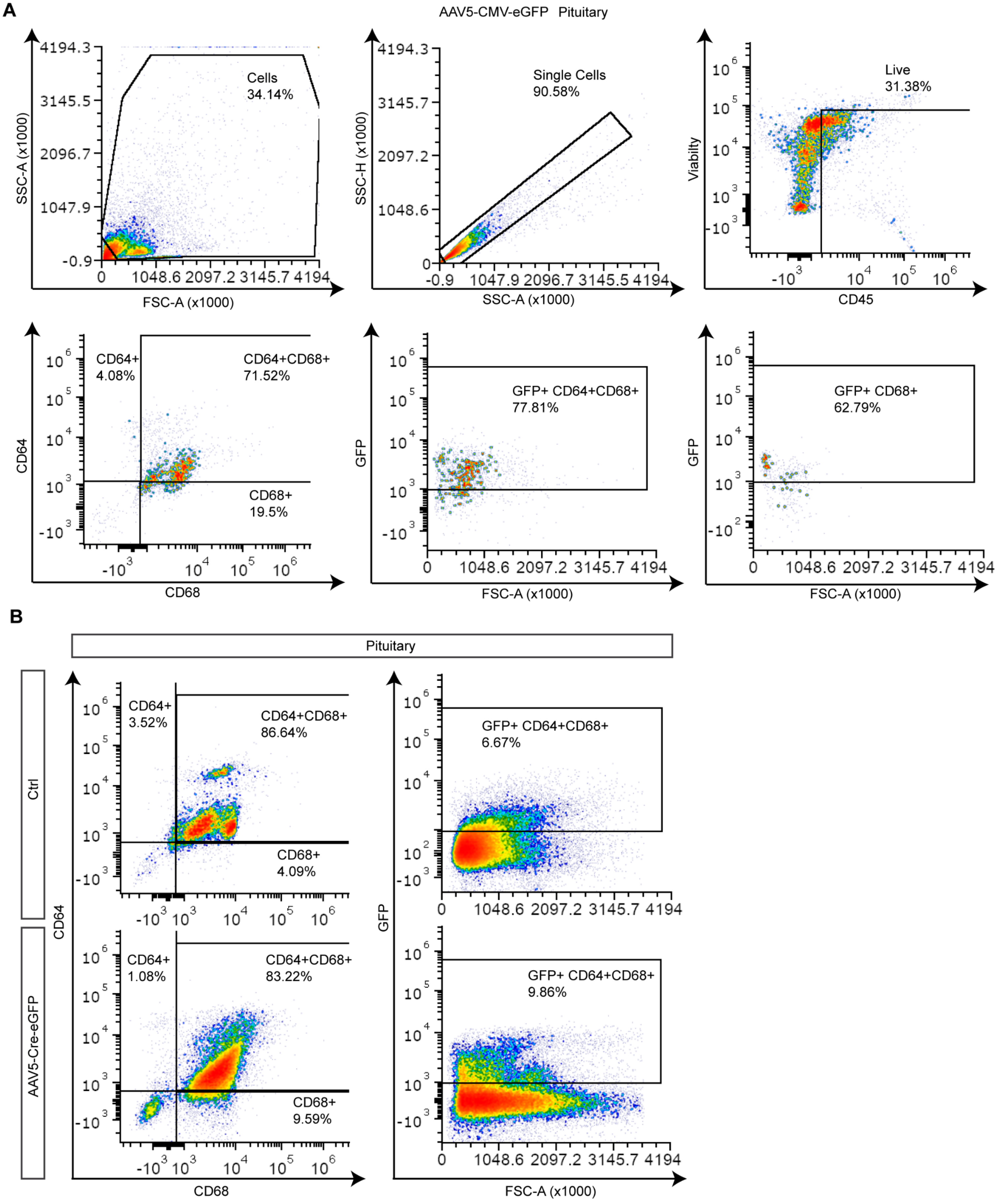


**Supplementary Figure 8.** (**A**) Flow cytometry gating strategy for pituitaries collected from C57BL/6J mice following retro-orbital injection of AAV5-CMV-eGFP. (**B**) Flow cytometry gating strategy for pituitaries collected from LysMCre mice following retro-orbital injection of AAV5-Cre-eGFP.

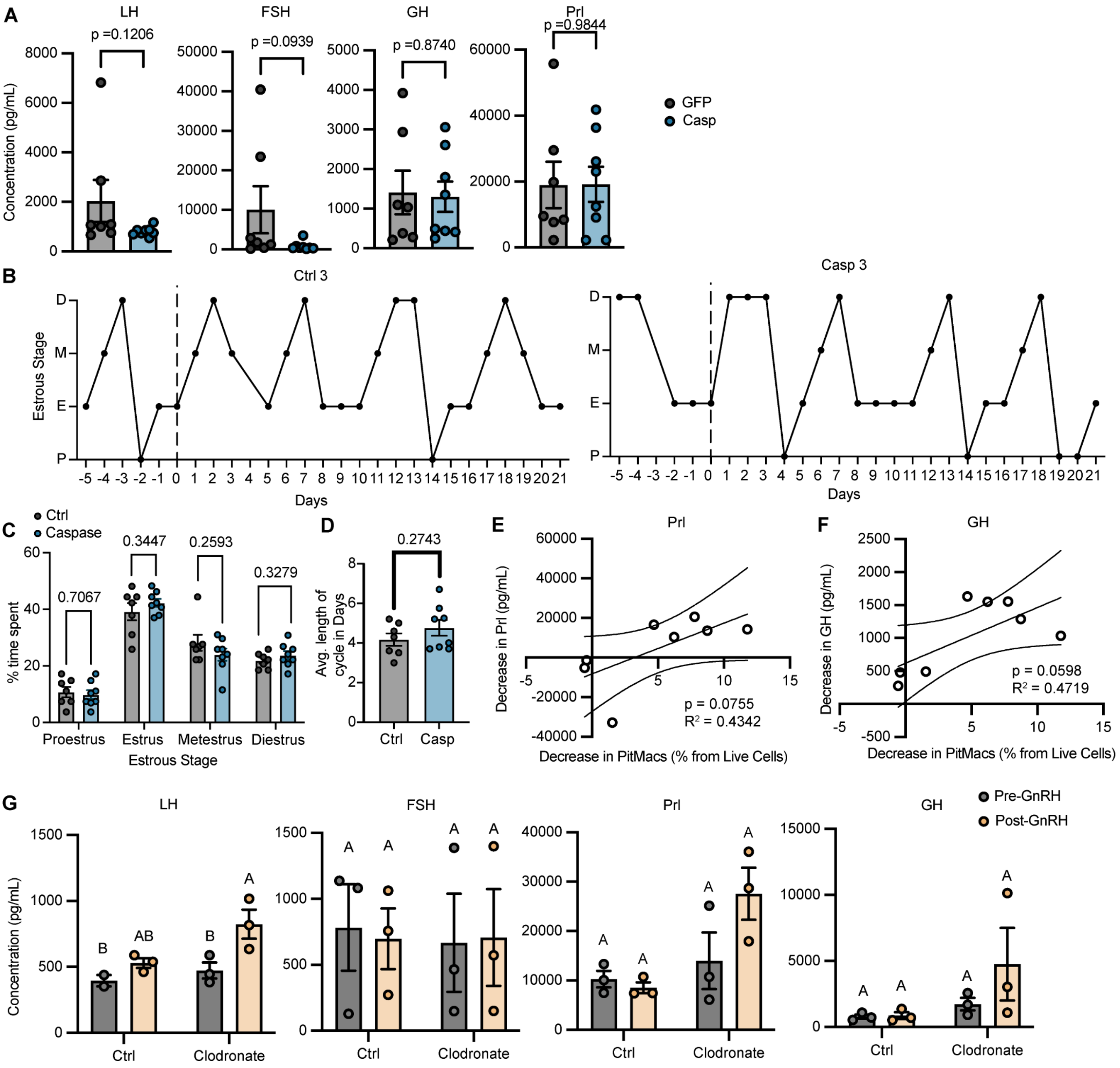


**Supplementary Figure 9.** (**A**) Serum LH and FSH concentrations (pg/mL) pre-GnRH challenge were measured by Luminex. Hormone concentrations and macrophage populations were analyzed following appropriate outlier exclusion tests, normality tests, and Box-Cox transformation. Mann-Whitney test was used for LH and FSH. Unpaired t-test was used for Prl and GH. *p = 0.01–0.05; **p = 0.001–0.01. (**B**) Estrous cycle staging data from 5 days prior to and 21 days following AAV-Cre-Casp injection, shown for representative mice from each treatment group. (**C**) Percentage of time spent in each estrous cycle stage in control and caspase-treated mice. (**D**) Average estrous cycle length in control and caspase-treated mice. (**E-F**) Pearson correlation analysis depicting the magnitude of decrease in serum Prl and GH levels (pg/mL) versus the magnitude of decrease in the PitMac population (% of live cells). Dashed lines represent 95% confidence bands of the best-fit line. (**G**) Serum LH and FSH concentrations measured by Luminex 48 hours following delivery of clodronate or control liposomes.

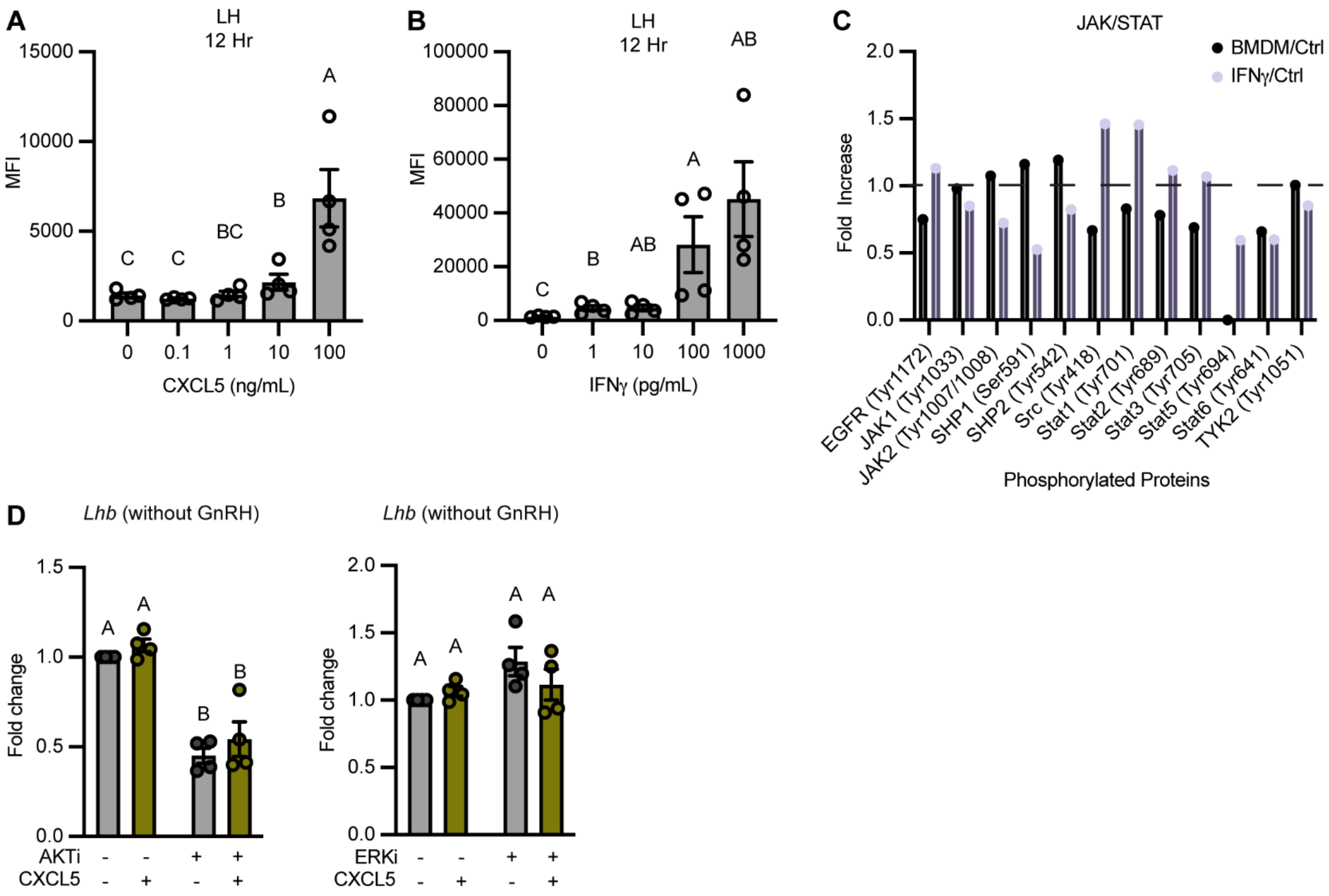


**Supplementary Figure 10.** (**A-B**) LβT2 cells were stimulated with increasing doses of CXCL5 (0.1,1,10, 100 ng/mL) (**B**) or IFN-γ (1,10,100,1000 pg/mL) (**B**). LH concentration in culture supernatants was measured by Luminex. Statistical comparisons were performed using Tukey's Honest Significant Difference (HSD) test following Box-Cox transformation; groups not sharing the same letter are significantly different. (**C**) Phospho-protein array analysis using a mouse JAK/STAT signaling pathway kit. LβT2 cells and BMDMs were treated with 100 pg/mL IFN-γ for 30 minutes. LβT2 cells were not serum starved prior to treatment.

### Supplementary Materials and Method

#### *In vivo* Macrophage Depletion by Clodronate Liposomes

Estrous stage was assessed 24 hours prior to injection in six 11-week-old female C57BL/6J mice (see Estrous Cycle Assessment). Clodronate or control liposomes (Encapsula NanoSciences, Cat. No. CLD-8901) were administered via retro-orbital injection. At 24 hours post-injection, estrous stages were reassessed and a GnRH challenge was performed (see GnRH Challenge). Pre-GnRH blood was collected via the retro-orbital sinus under isoflurane anesthesia, and post-GnRH blood was collected from trunk blood following euthanasia. Pituitaries and spleens were harvested for flow cytometric analysis of macrophage populations.

#### Western Blot

Pituitaries lysates were used from a previous experiment *(28)* **and probed for CD45.**
